# Integrating Biotic Interactions In Niche Analyses Unravels Patterns Of Community Composition in Clownfishes

**DOI:** 10.1101/2023.03.30.534900

**Authors:** Alberto García Jiménez, Antoine Guisan, Olivier Broennimann, Théo Gaboriau, Nicolas Salamin

## Abstract

Biotic interactions shape the ecology of species and communities, yet their integration into ecological niche modeling methods remains challenging. Despite being a central topic of research for the past decade, the impact of biotic interactions on species distributions and community composition is often overlooked. Mutualistic systems offer ideal case studies for examining the effects of biotic I nteractions on species niches and community dynamics. This study presents a novel approach to incorporating mutualistic interactions into niche modeling, using the clownfish-sea anemone system. By adapting existing niche quantification frameworks, we developed a method to estimate the partial effects of known interactions and refine ecological niche estimates. This approach allows for a more comprehensive understanding of how mutualistic relationships influence species distributions and community assembly patterns. We also used mutualistic information to investigate the resource-use overlap, identitying patterns of competition within clownfish communities. Our results reveal significant deviations in niche estimates when biotic interactions are considered, particularly for specialist species. Host partitioning among clownfish species reduces resource-use overlap, facilitating coexistence in species-rich habitats and highlighting mutualism’s role in promoting and maintaining diversity. We uncover complex dynamics in resource-use overlap among clownfish species, influenced by factors such as species richness, ecological niche overlap, and host specialization. Specialist-generalist interactions strike an optimal balance, supporting high species richness while minimizing competition. These insights enhance our understanding of clownfish biodiversity patterns, demonstrating how diverse mutualistic strategies contribute to diversity build-up and mitigate competitive exclusion in saturated communities. The analytical framework presented has broad applications beyond the clownfish-sea anemone system, potentially extending to a broader range of interactions. It enables a more comprehensive understanding of biodiversity maintenance in complex ecosystems and constitutes a valuable tool for conservation planning and ecosystem management.

## Introduction

Diverse types of biotic interactions govern the interconnection between species in nature. Among them, mutualism, a relationship in which different species benefit from each other, has attracted the attention of biologists and the public at large. Mutualism plays a substantial role in evolutionary processes. It has impacted major evolutionary transitions like the origin of the eukaryotic cell or the colonization of land by symbiotic plants, and contributes to the increase of biodiversity (Bastolla *et al*. 2009) by enhancing the survival and success of interacting species (Benton 2009). It also has an important ecological impact by favouring ecosystem stability, and facilitating species dispersal and resilience (Hale *et al*. 2020; Le Roux *et al*. 2020), impacting species distributions (Pellissier *et al*. 2013; Schleuning *et al*. 2015; Marjakangas *et al*. 2020).

Mutualistic interactions present a gradient of intensity from full generalists to exclusive specialists, leading to diverse ecological and evolutionary dynamics (Bascompte & Jordano 2007; Sverdrup-Thygeson *et al*. 2017; Gracia-Lázaro *et al*. 2018). Generalist mutualism promotes ecosystem resilience through flexible responses to environmental changes (Maia *et al*. 2021). In contrast, specialist mutualism can drive co-evolution and lead to highly adapted species pairs (Cook & Rasplus 2003). Generalist-specialist transitions can foster diversification through ecological speciation (Lunau 2004), where reproductive isolation evolves as a by-product of adaptation to different ecological niches (Chomicki *et al*. 2019; Frachon *et al*. 2023). As species adapt to specific mutualistic partners, they may undergo niche partitioning, reducing niche overlap to avoid competitive exclusion (Salas-Lopez *et al*. 2022), allowing closely related species to coexist in the same ecosystem (Schoener 1974). Plant-pollinator systems exemplify this process, with related plant species evolving to attract different pollinators or flower at different times, reducing pollination competition (Van der Niet *et al*. 2014). In the marine environment, the clownfish (Amphiprioninae) mutualism with host sea anemones might have triggered their rapid diversification through ecological speciation, a process known as adaptive radiation (Litsios *et al*. 2012).

The evolutionary success of this group of 30 reef fish species, is attributed to their unique mutualistic associations with sea anemones in the Indo-Pacific Ocean. The mutualism significantly enhances clownfish survival and reproductive success (Lubbock 1980; Fautin 1991), with each species developing specific associations with up to ten sea anemone species, resulting in both generalist and specialist behaviors. Despite their similar ecological characteristics (habitat, diet and social structure), clownfishes exhibit higher levels of coexistence in species-rich locations (Camp 2016; Elliott & Mariscal 2001), contrary to expectations of increased competition. We hypothesize that mutualistic interactions with sea anemones facilitate niche partitioning among clownfish species, reducing interspecific competition and enabling coexistence. This mechanism would provide ecological support for the adaptive radiation hypothesis, demonstrating mutualism’ role in promoting clownfish diversity. However, conclusive evidence of niche partitioning among clownfish species is lacking, hampering the testing of adaptive radiation hypotheses and challenging our understanding of clownfish diversification.

Furthermore, the interplay between generalist and specialist strategies likely influences clownfish distributions and community assembly (Chesson 2020; Bartholomew *et al*. 2022; Wandrag *et al*. 2022). Communities dominated by specialist-specialist interactions are expected to show limitations in species richness due to increased competition for specific anemone hosts. Conversely, communities with more generalist interactions could support higher diversity by reducing direct competition and allowing for more efficient resource utilization. Investigating this interplay could reveal spatial patterns linked to species richness, elucidating clownfish ecological roles, and deepening our understanding of how mutualistic interactions shape community structure, species coexistence, and biodiversity patterns.

To test these hypotheses, we need to estimate the realized environmental niches of clownfish species, while accounting for the effect of explicit mutualistic associations. However, current approaches quantifying ecological niches and modeling species distributions involve techniques like principal component analysis (PCA; Broennimann *et al*. 2012) and niche-based spatial modelling of species distributions (SDMs; see Guisan *et al*. 2014, Valavi *et al*. 2021; Norberg *et al*. 2019), do not directly incorporate biotic interactions. This is despite the demonstrated improvement in SDM predictions that the integration of biotic interactions can bring (Wisz *et al*. 2013; Early & Keith 2019; Kass *et al*. 2020; Jenkins *et al*. 2020) and the recurrent calls to develop SDMs that can integrate these interactions (see Soberón 2010; Boulangeat *et al*. 2012; de Araujo *et al*. 2013; Leach *et al*. 2016; D’Amen *et al*. 2018; but see Konig *et al*. 2021). Consequently, the role of biotic interactions is frequently overlooked (Anderson 2017; Palacio & Girini 2018), which can lead to misinterpretations of ecological niches and species distributions (Zurell *et al*. 2020; Moullec *et al*. 2022). In our case, such misinterpretations would hinder our capacity to fully understand the significant role that mutualism plays in shaping the ecological niche of clownfishes.

Here we built on an existing environmental niche quantification approach and implemented a novel method to reduce the biotic uncertainty, estimate the partial effects of known interactions, and accordingly refine the estimate of the ecological niche. To capture the mutualistic dependence of clownfishes on sea anemones and understand their effects on clownfishes niche, we adapted the ‘COUE’ framework (Guisan *et al*. 2014) by defining new metrics describing the effect of biotic interactions on niche quantification (Fig. 1). We compared the impact of biotic interactions on niche quantification between generalist and specialist clownfishes, hypothesizing that sea anemone associations significantly affect clownfish niches and distributions, with a greater effect on specialists than on generalists. Furthermore, we anticipate that incorporating mutualistic interactions as a limited resource into the quantification of resource overlap will reflect current competition dynamics among clownfishes, as well as their relationship to species richness and community assembly patterns.

**Fig. 1:**
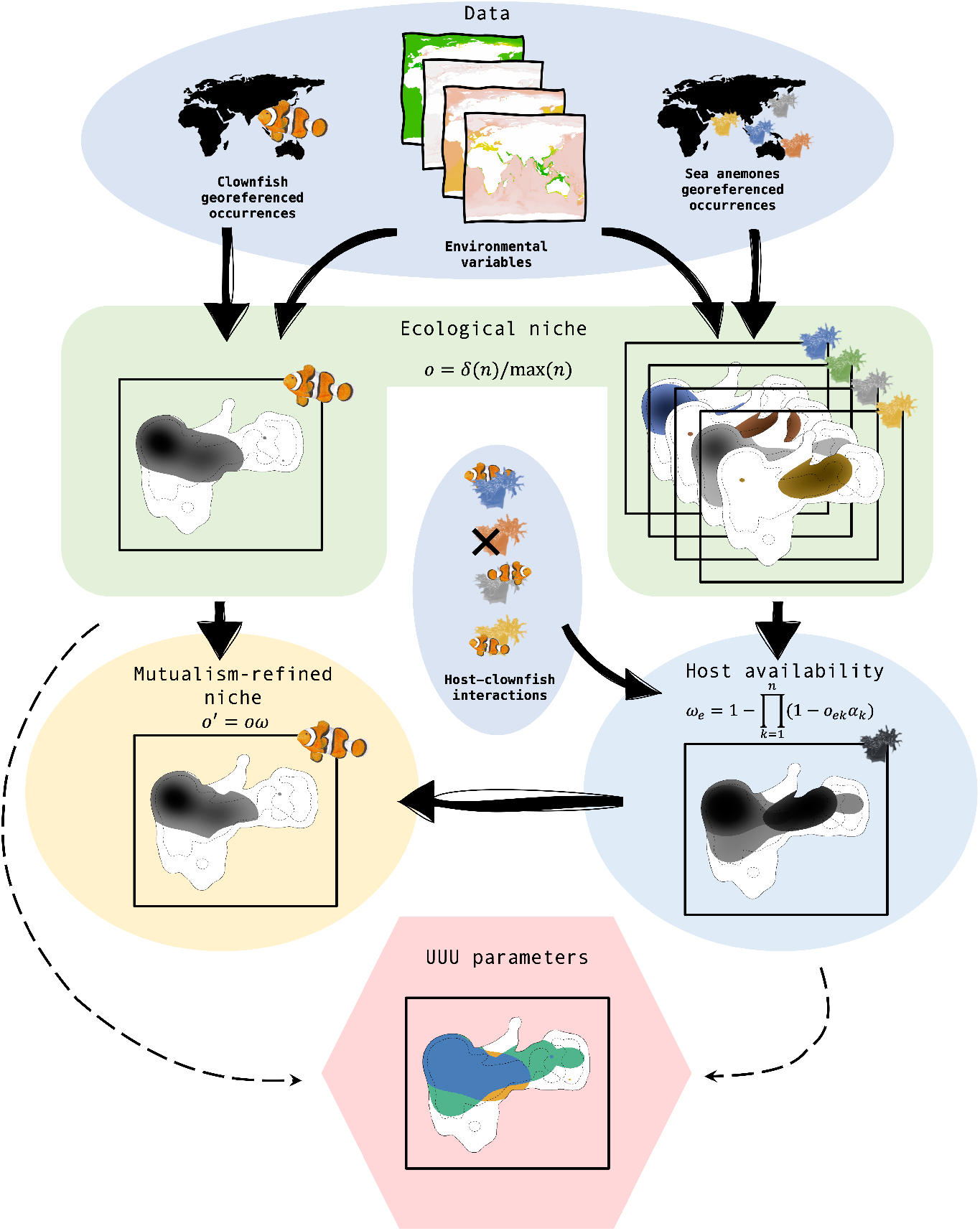
Scheme of the proposed framework to assess the effect of biotic interactions and whether differential mutualistic behaviors may produce biases in the ecological niche estimation. For each clownfish species, georeferenced occurrences were collected, and ecological niche was estimated from the environmental space created using the selected environmental variables. Additionally, georeferenced occurrences of all sea anemones species were collected to infer their ecological niches following the same procedure as for the clownfish. Hosts ecological niches were combined into a single multi-hosts ‘niche’ using the interaction vector following the provided formula to create an envelope of host availability (*ω*). Finally, we constrained the clownfish ecological niche (*o*) by the host availability (*ω*) to obtain a mutualism-refined ecological niche (*o*^*′*^). Comparisons between the host availability envelope and the estimated clownfish ecological niche (dashed lines) provided the UUU parameters, determining Unavailable environments (environmentally suitable but unavailable due to lack of host availability), Used environments as they were both suitable and available, and Unoccupied environments as those that were available but not suitable for the clownfish.

## Material & Methods

### Data Collection

Clownfishes inhabit the Indo-Pacific Ocean, from the East coast of Africa to Polynesia in the Pacific, and from the coast of Japan to the South of Australia, while sea anemones have a broader, worldwide distribution (Fautin & Allen 1992). We collected 1,636 occurrences of ten host sea anemone species (mean: 163.6; min: 68; max: 335) and 4,258 occurrences of 30 clownfish species (mean: 146.8; min: 2; max: 860) from RLS, GBIF, OBIS and Hexacoral databases (Atlas of Living Australia 2017; GBIF.org 2018; OBIS 2017; Fautin 2008 respectively). Datasets were filtered to remove duplicates and misplaced or misidentified occurrences. After filtering, two clownfish species (Amphiprion pacificus and thiellei) with fewer than five occurrences were excluded.

Environmental data from GMED (Basher *et al*. 2018) and Bio-Oracle (Tyberghein *et al*. 2012; Assis *et al*. 2018) were obtained at 0.083° resolution, representing approximately 9,2 km near the equator, covering physical, chemical, and biological factors. The 53 environmental variables collected were restricted to shallow reefs and the epipelagic zone above 50m depth, using the UNEP-WCMC warm water coral reef map (UNEP-WCMC 2018). After removing variables with excessive missing data and discarding highly correlated variables (Pearson’s correlation > 0.8), we retained eight environmental variables: mean current velocity, mean salinity, mean temperature, mean nitrate concentration, nitrate concentration range, mean chlorophyll concentration, dissolved oxygen concentration range, and mean phytoplankton concentration.

### Quantifying ecological niches

We constructed a global environmental space using the first two principal components of a scaled PCA based on the eight selected environmental variables, which explained 39% and 22% of the total variance, respectively. We used the *ecospat* R package (Broennimann *et al*. 2012) to estimate the occurrence density o for both clownfishes and sea anemones in a two-dimensional grid of 100 by 100 cells:

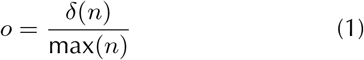

where δ(*n*) is the 2D kernel density estimation of the number of occurrences on the defined environmental space, and max(*n*) is the maximum number of occurrences in any grid cell of the environmental space. The occurrence density *o* ranges from 0 for environments where the species is not observed to 1 where it is most observed. It represents the species ecological niche (i.e. realized environmental niche) and can be used as a proxy of the probability of occurrence in a given environment (Drake & Richards 2018)..

### Refining the ecological niche using mutualistic interactions

We compiled the species-specific mutualistic interactions between sea anemones and clownfishes from scientific literature (Fautin 1985, 1991; Godwin & Fautin 1992; Ollerton *et al*. 2007; Ricciardi *et al*. 2010; Litsios *et al*. 2012) and reputable online sources (https://amphiprionology.wordpress.com, www.fishbase.org, https://reeflifesurvey.com) to construct an association matrix 𝒜 where each clownfish species *s* was represented by a vector of interactions (Fig. 1):

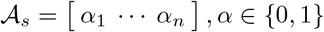

with *n* being the number of sea anemone species (here 10 in total). Given the obligate nature of the clownfish-anemone mutualism, we assumed that environments unsuitable for host anemones would be unavailable to clownfish, resulting in *α* values of either 0 or 1 in our case (see Supplementary Material & Methods for potential extensions). Environmental availability for clownfish would thus depend on two factors: i) the association between a present host species and the clownfish, as defined by matrix 𝒜, and ii) the occurrence density of sea anemone species in the environment. The host availability *ω* was estimated as

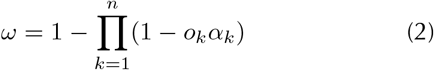

where *α*_*k*_ is the association between a clownfish and its host anemone *k* (derived from the matrix 𝒜), while _*k*_ is the occurrence density of the host *k. ω* values range from 0 to 1 and represent the availability score across the two-dimensional environmental space where a suitable host is present. Then, the refined occurrence density *o*’ of the focal clownfish species given the mutualistic associations was

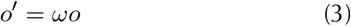

where *o* is the occurrence density of the focal clownfish species, and *ω* is the host availability across the environmental space.

Given clownfishes’ limited dispersal capacity (Jones *et al*. 2005; Almany *et al*. 2017) and local adaptation of both clownfishes (Huyghe & Kochzius 2015; Clark *et al*. 2021; Ducret *et al*. 2022) and sea anemones (Sachkova *et al*. 2020; Will *et al*. 2021; Prakash *et al*. 2021), we anticipated regional ecological variations. We divided the study area into 27 regions across 5 marine realms based on MEOW (Spalding *et al*. 2007; Fig. S1). We estimated the ecological niche *o* for both clownfish and sea anemones, and the mutualism-refined niche *o*’ for clownfishes, at regional scales by assessing species niches within each marine region’s environmental space subset. Main analyses were conducted at a global scale and without environmental variable selection to test the robustness of our findings across spatial scales and variable inclusion (Fig. S2 and S3). Additionally, we projected the occurrence densities of each estimated niche into the geographical space for spatial representation of the results (Supplementary Material & Methods). We also conducted sensitivity analyses to evaluate the effect of the association matrix in our framework (Fig. S4 and Supplementary Material & Methods).

### Effect of mutualistic interactions on species niches

We compared the clownfishes niche estimates with and without accounting for the sea anemones hosts occurrences (ecological niche *o* vs. mutualism-refined niche *o*’) to understand the effect of implementing explicit mutualistic interactions in niche estimations. We measured different characteristics of niche space by adapting the ‘COUE’ framework (Guisan *et al*. 2014). In particular, we quantified changes in species’ realized niches by defining three categories of occupied environmental space, accounting for the influence of sea anemone hosts. With *N* representing the number of cells in the environmental grid, we defined ‘Unavailable’ as the proportion of environmental space not occupied by the hosts of the given clownfish species and thus not available for the clownfish, calculated as *N*_(*o>*0_ & _*o*’=0)_/*N*_(*o>*0)_, ‘Used’ as the proportion of environmental space occupied by the given clownfish species, calculated as *N*_(*o>*0_ & _*o*’*>*0)_/*N*_(*o>*0)_, and ‘Unoccupied’ as the proportion of environmental space inhabited by any host of the given clownfish species but not by the clownfish, calculated as *N*_(*o*’*>*0)_/*N*_(*ω>*0)_. To evaluate how the generalist-specialist spectrum affects ecological niche estimates incorporating mutualistic data, we classified species based on host associations. Following Ollerton *et al*. (2007) and Litsios *et al*. (2014), we defined generalists as species interacting with three or more hosts, and specialists as those with fewer than three hosts.

### Effect of mutualistic interactions on species niche overlaps

We examined how clownfish mutualistic interactions influence niche overlap among clownfish species. Using Schoener’s *D*, we calculated pairwise ecological niche overlap between species, both with and without considering host anemone associations. Ecological and mutualism-refined niche overlap inadequately represents actual competition in clownfish, which is primarily driven by host use. To estimate resource-use competition more accurately, we developed a multi-layered approach. We replicated the environmental space into layers corresponding to each host, projected species ecological niches onto these layers, and constrained them by host suitability (skipping equation 2 and taking each host’s occurrence density as *ω*). We then averaged layer-specific overlap values to obtain a single overlap measure for each species pair (Fig. S5). This method accounts for the host-specific nature of clownfish competition and provides a more realistic estimate of resource-use overlap. By integrating host specificity and environmental preferences, it captures the interplay between habitat requirements and resource competition in the clownfish-sea anemone mutualism system. We compared overlap estimates (ecological niche, mutualism-refined niche, and resource-use) and examined differences based on species pair specialization levels (generalist-generalist, generalist-specialist, and specialist-specialist).

### Spatial patterns of clownfish interspecific niche overlap

We examined the relationship between species richness and both ecological niche and resource-use overlap, as well as their geographical patterns. Species richness was defined as the number of species predicted at a location, while overlap intensity was calculated as the average pairwise overlaps of all species at a site. Using spatial generalized linear mixed models (GLMMs) with the *spaMM* R package (Rousset & Ferdy 2014), we assessed the effect of species richness on both overlap estimates (ecological niche and resource-use), accounting for spatial autocorrelation (Fig. S6, S7 and Table S1). We then analyzed subsets of ecological niche and resource-use overlap for generalist-generalist, specialist-generalist, and specialist-specialist interactions, estimating species numbers for each interaction type. Finally, we conducted spatial GLMMs for each subset to determine if niche overlap patterns varied by interaction type.

## Results

### Effect of mutualistic interactions on species niches

Incorporating mutualistic information significantly altered ecological niche estimates in 60% of the 108 regional niche subsets (Table S2). Across all clownfish species and regions, an average of 66% of a species ecological niche remained used after integrating mutualistic Information 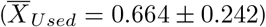. Approximately one-sixth was unavailable due to unsuitable host environments 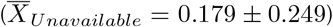, while a similar proportion was suitable for hosts but outside the clownfish ecological niche 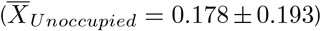. UUU proportions however, exhibited considerable variability among species (Fig. 2) and across regions (Fig. S8).

**Fig. 2:**
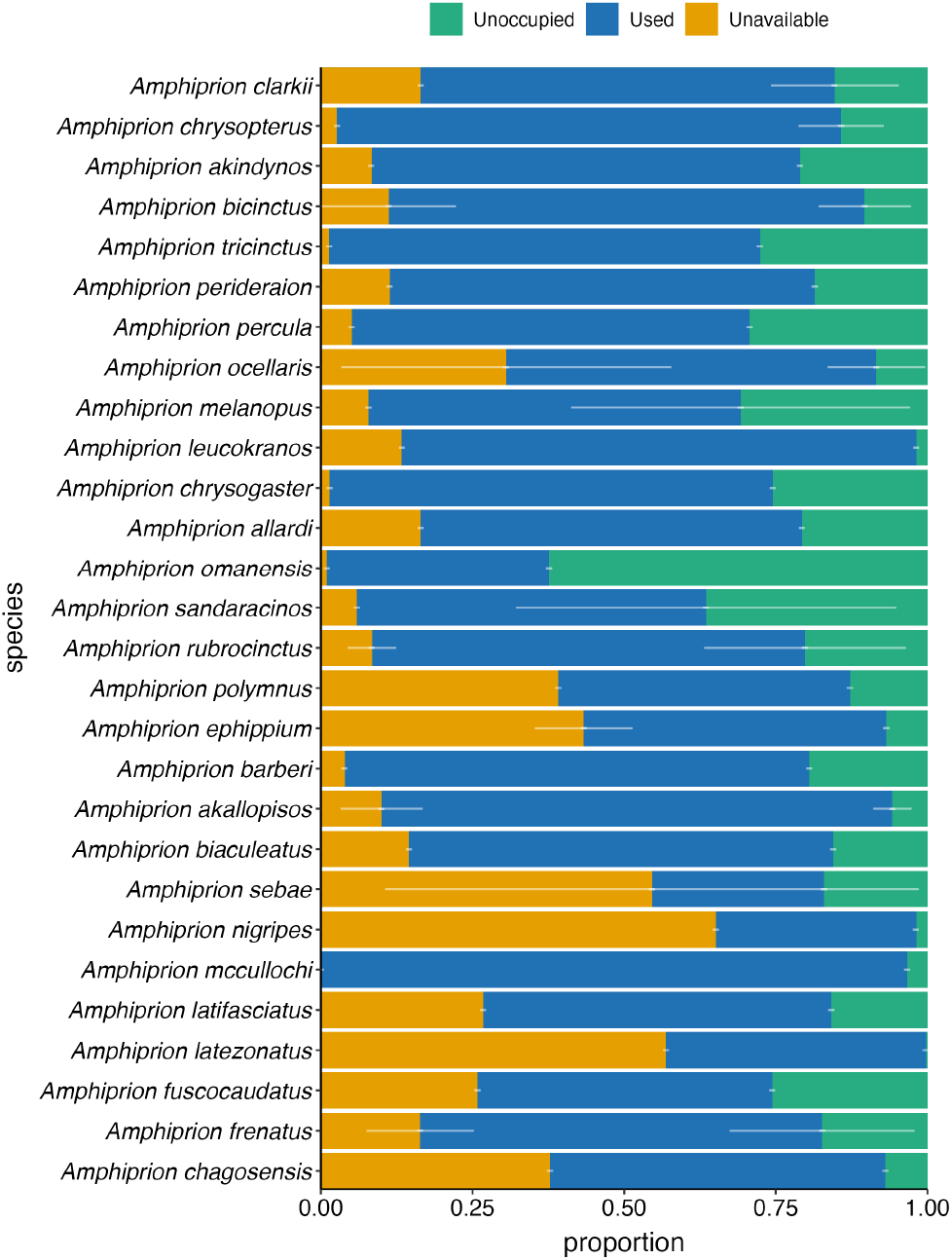
Stacked bar plot showing the averaged UUU proportions per clownfish species among regions. Colours represent the different UUU parameters specified on the legend on top. White vertical lines represent the standard deviation across regions.

Specialists and generalists showed significant variations in UUU proportions (Fig. 3 and Table S3). Specialists had higher Unavailable (*H* = 23.061, *df* = 1, *p <* 0.001) and lower Used (*H* = 17.129, *df* = 1, *p <* 0.001) proportions in their ecological niches compared to generalists, while no significant difference was detected for Unoccupied proportions (*H* = 2.1735, *df* = 1, *p* = 0.14). The stronger effects on specialist clownfish species Unavailable and Used proportions were evident through higher niche dissimilarity (*H* = 39.987, *df* = 1, *p <* 0.001), greater centroid shift (*H* = 20.681, *df* = 1, *p <* 0.001), and larger environmental shift (*H* = 17.704, *df* = 1, *p <* 0.001) when compared to generalists (Fig. S9). These UUU patterns were consistent at the spatial level (Fig. S10)

**Fig. 3:**
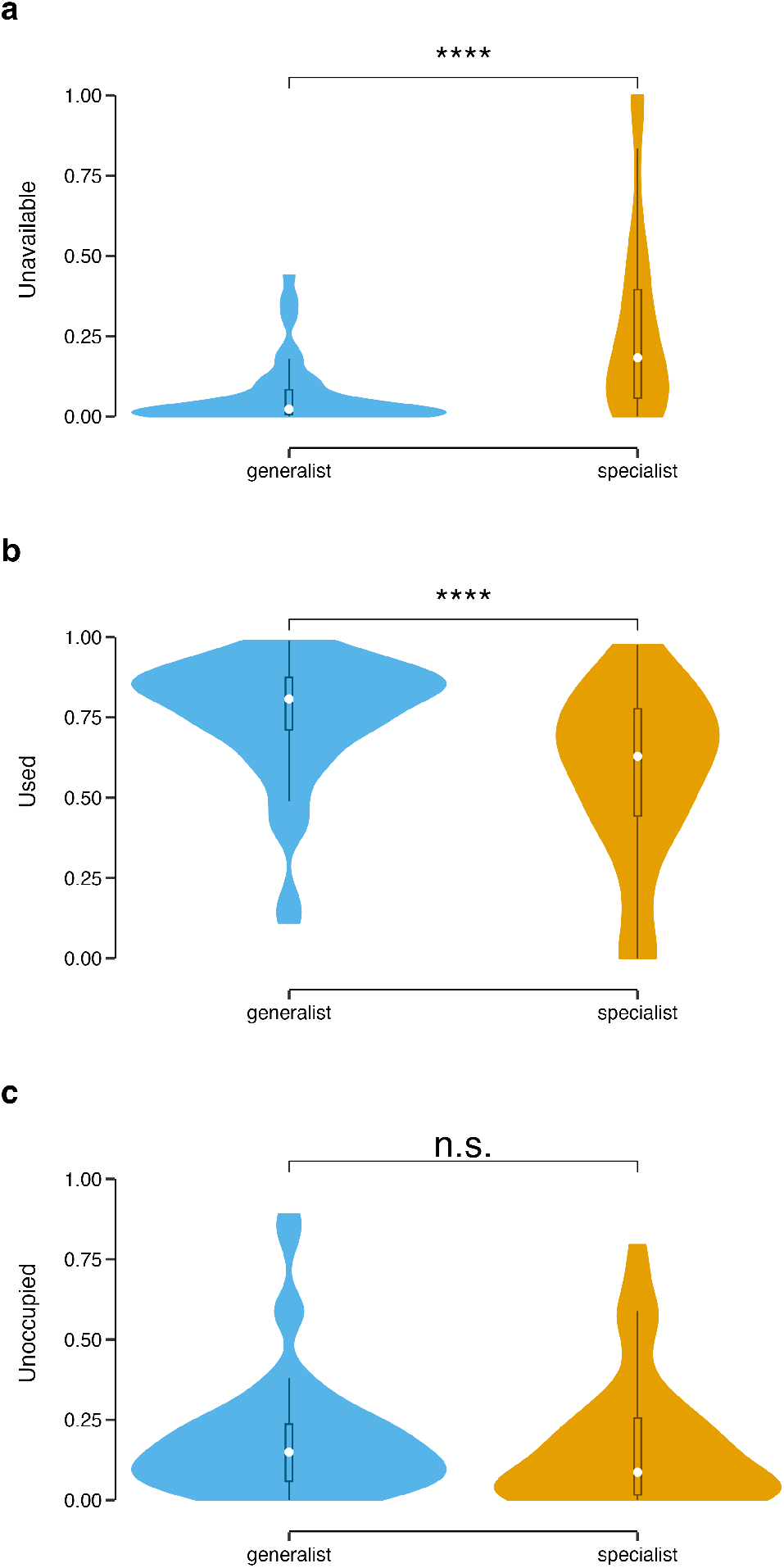
Comparisons between generalists and specialists on the proportions of Unavailable (a), Used (b), and Unoccupied (c) proportions of the niche, adapted from the ‘COUE’ framework (Guisan *et al*. 2014). Violin plots show the distribution of the data. Statistical significance is represented following the legend: no significant (n.s.); *p* < 0.05 (*); *p* < 0.01 (**); *p* < 0.001 (***); *p* < 0.0001 (****).

### Effect of mutualistic interactions on species niche overlaps

Clownfish ecological niche overlap, measured by Schoener’s *D* statistic, was high (median *D*_*ecological*_ = 0.741; *IQR* = 0.199), indicating highly similar environmental suitability. Accounting for host associations slightly but significant (*V* = 14, 723; *df* = 1; *p* = 0.015) increased niche overlap (median *D*_*mutualism*−*refined*_ = 0.762; *IQR* = 0.220) compared to ecological niche overlap. However, resource-use overlap was substantially lower (median *D*_*resource*−*use*_ = 0.232; *IQR* = 0.304) and significantly different from ecological and mutualismrefined niche overlap (*V* = 35, 327; *df* = 1; *p <* 0.001 and *V* = 33, 650; *df* = 1; *p <* 0.001, respectively; Fig. 4).

**Fig. 4:**
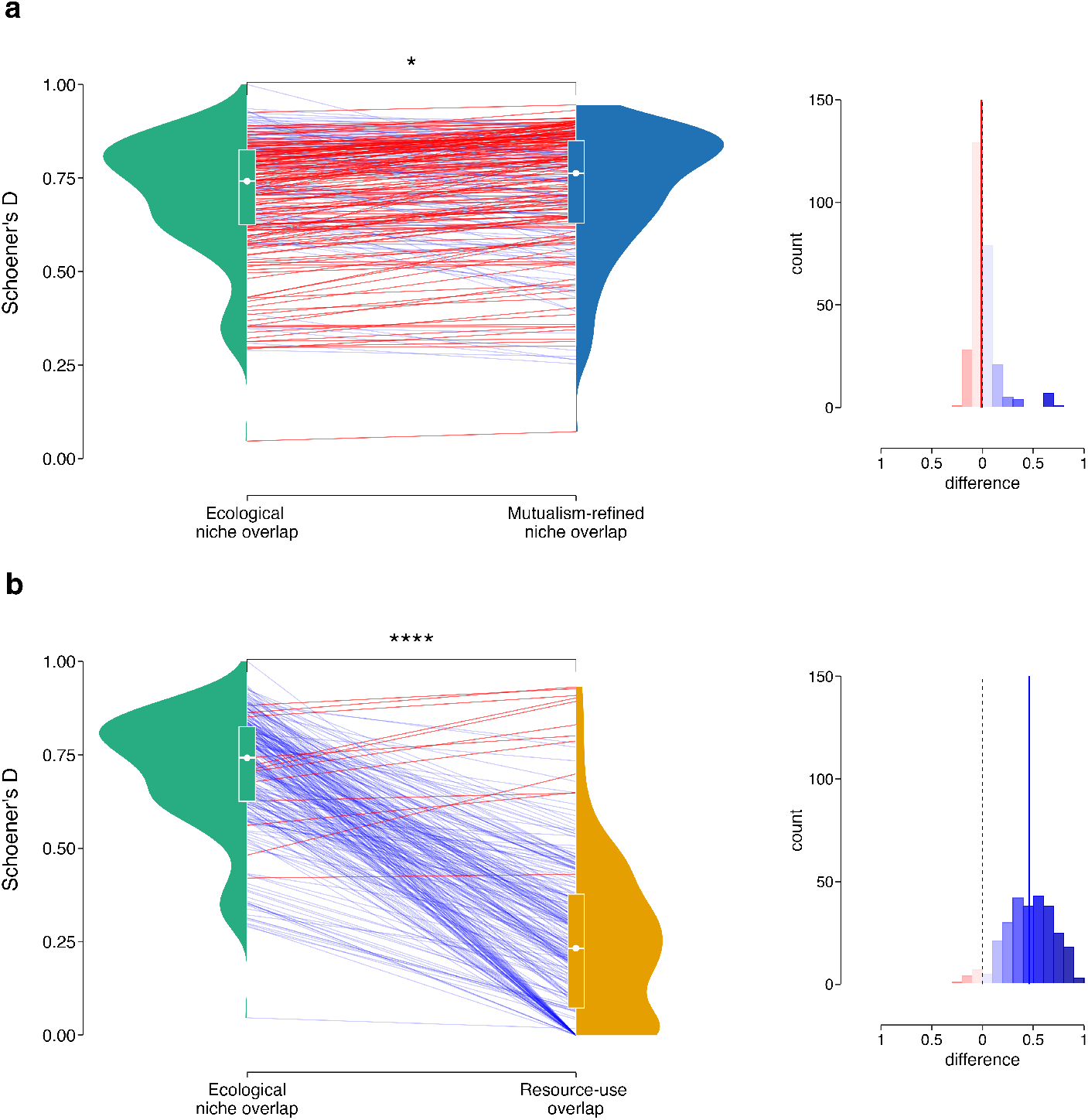
Pairwise species comparison of between ecological and mutualism-refined niche overlaps (a), and ecological niche and resource-use overlaps (b). Colored lines in the main plots (left) represent individual pairwise comparisons across categories, with blue lines indicating a decrease in overlap intensity and red lines showing an increase. Boxplots illustrate the overall intensity of niche overlap at each niche level, while violin plots show the distribution of pairwise overlaps for each category. Histograms on the right display the distribution of differences between pairwise overlaps, where blue bars indicate decreased overlap and red bars indicate increased overlap compared to ecological niche overlap. Statistical significance is represented following the legend: no significant (n.s.); *p* < 0.05 (*); *p* < 0.01 (**); *p* < 0.001 (***); *p* < 0.0001 (****).

*D*_*ecological*_ showed no significant differences among specialist-specialist, specialist-generalist, generalist-generalist interactions (*H* = 4.193; *df* = 2; *p* = 0.122). However, significant differences emerged after refining the ecological niche with mutualistic information (*H* = 6.7509; *df* = 1; *p* = 0.034), with stronger differences in resource-use overlap (*H* = 71.623; *df* = 1; *p <* 0.001). In specialist-specialist pairs, 15 comparisons showed zero resource-use overlap due to not shared hosts, while the remaining 13 pairs exhibited high overlap (median *D*_*resource*−*use*_ = 0.456; *IQR* = 0.374). Overlap decreased significantly in generalist-generalist (median *D*_*resource*−*use*_ = 0.351; *IQR* = 0.212) and generalist-specialist interactions (median *D*_*resource*−*use*_ = 0.201; *IQR* = 0.172), with the latter, being the most common among co-occurring clownfishes and exhibiting the lowest resource-use overlap (Fig. S11).

### Geographical patterns of niche overlap

The Eastern and Western Coral Triangle regions showed the highest species richness and number of potential interactions (Fig 5a). High environmental suitability overlap was found in Pacific, Western and Central Indian Oceans regions for both ecological and mutualism-refined niches projections (Fig 5b; ecological niche projections shown). Resource-use overlap was more pronounced in the Tropical North-western Pacific and Somali/Arabian sea (Fig 5c). Generally, resource-use overlap was lower than environmental suitability overlap for within regions (Fig 5d), with the largest disparities in Temperate and Tropical Pacific regions, Northeast Australian Shelf, Coral Triangle, and Western Indian Ocean.

**Fig. 5:**
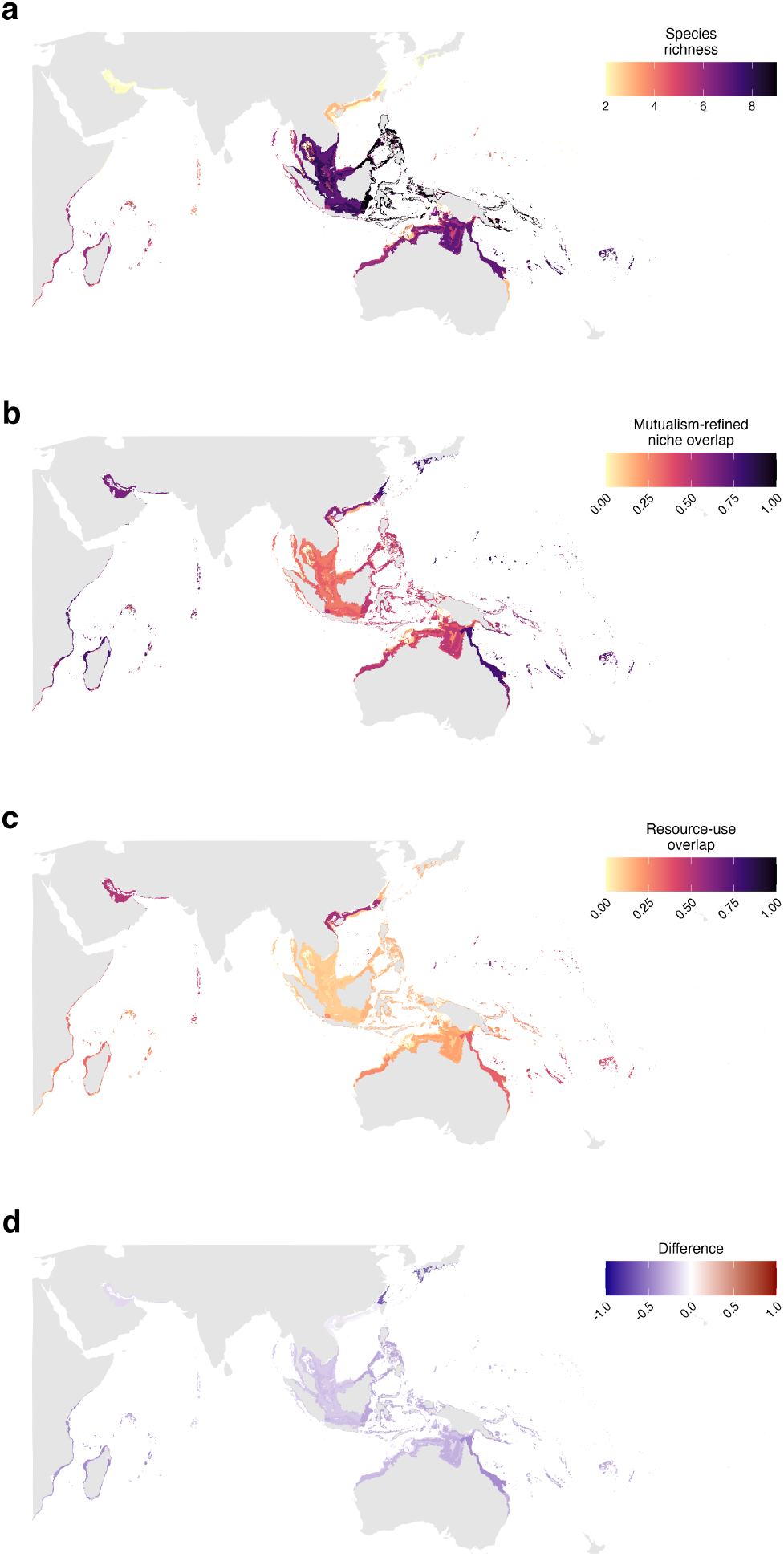
a) Estimated number of clownfish species occurring per location. b) Averaged ecological niche overlap among all co-occurring species per location. c) Average resource-use overlap among of all co-occurring species per location. d) Difference between ecological niche overlap and resource-use overlap computed as D_resource-use_ - D_ecological_. Negative values represent higher ecological niche overlap than resource-use overlap and positive values represent higher resource-use niche overlap than ecological niche overlap.

Resource-use overlap correlated positively with ecological overlaps across regions (*r* = 0.800, *p <* 0.001) and negatively with species richness (*r* =−0.287, *p <* 0.001). GLMMs revealed species richness negatively associated with resource-use overlap (*β* = − 0.013; 95%*CI*[− 0.015, − 0.017]), while environmental suitability overlap (*β* = 0.668; 95%*CI*[0.646, 0.689]) showed a positive association. Their interaction had a negative impact on resource-use overlap (*β* = 0.015; 95%*CI*[− 0.011, 0.019]; Fig. S12 and Table S4). Analyzing interaction subsets (generalist-generalist, specialist-generalist, and specialist-specialist), resource-use overlap among generalist-generalist interactions showed positively associated with generalist species richness (*β* = 0.013; 95%*CI*[0.009, 0.017]) and environmental suitability overlap (*β* = 0.415; 95%*CI*[0.400, 0.430]). Specialist-generalist and specialist-specialist interactions exhibited similar effects as the overall model, with varying magnitudes of association between environmental suitability and resource-use overlap (Table S4).

## Discussion

Our study introduces a novel approach to incorporate mutualistic interactions in the estimation of ecological niches. Using clownfishes as a case study, we explored how species-specific mutualistic interactions with sea anemones influence the ecological niche. This allowed us to assess niche overlaps among clownfishes based on resource use, shedding light on competition patterns within clownfish communities. We then evaluated the role of mutualistic behaviour and host-partitioning in resource competing communities.

### Effect of mutualistic interactions in clownfish species niches

Our study revealed significant misalignment between clownfish and host anemone ecological niches (Fig 2). Despite their obligate mutualism, we found only about 60% overlap between clownfish and host niches, contrary to our expectation of nearly-identical environmental suitability. This discrepancy exceeds that observed in non-dependent interactions of other organisms (e.g. Arumoogum *et al*. 2023). While some mismatch could be attributed to data limitations and methodological constraints, the extent of disagreement was unexpected given the nature of their relationship.

Comparing ecological niches (based on environmental factors) with mutualism-refined niches revealed deviations due to the presence or absence of suitable host anemones. These disparities can introduce potential biases into spatial models, especially for specialists species heavily constrained by biotic interactions (Meineri *et al*. 2012). Our approach incorporates explicit information about mutualistic interactions, refining ecological niche estimations and enabling detailed examination of environmental suitability discrepancies between clownfish and their host anemones. Spatial projections of the UUU parameters showed ‘Used’ areas (suitable for both clownfish and host anemones) as predominant in species distributions (Fig. S13 and S14), indicating a strong agreement in environmental suitability. ‘Unavailable’ (suitable for clownfish only) and ‘Unoccupied’ (suitable for hosts only) areas represent less common environments, highlighting suitability mismatches between mutualistic partners (Fig. S15). The strong agreement between clownfish and host anemones ecological niches in predominant environments suggests an ecologically stable clownfish-sea anemone interaction, likely maximizing habitat occupancy in a convergent manner. This pattern may indicate co-evolution, with both partners adapting to similar environmental conditions, driven by their symbiotic benefits.

Regions with high ‘Unavailable’ proportions due to host absence may indicate ecologically unstable or fluctuating populations, potentially representing sink populations. These areas show inconsistent observations of both host anemones and clownfish. Such pattern could indicate non-self-sustaining populations relying on immigration from stable source populations. However, these sink populations might foster new ecological adaptations (Peniston *et al*. 2019). In source-sink dynamics, sink habitats can play crucial roles in adaptation by harboring increased genetic variation and experience reduced competition, allowing for survival of novel phenotypes (Holt, 1996; Kawecki, 2008). Temporal variation in harsh sink environments can facilitate niche evolution, with favorable periods allowing population growth and fixation of beneficial mutations (Lenormand, 2002; Holt *et al*. 2003). In clownfish-anemone systems, populations in low host availability areas might evolve wider environmental tolerances, utilize alternative hosts, or enhance dispersal capabilities. Long-term population genetic studies are needed to fully understand adaptive processes of clownfishes in marginal habitats.

‘Unoccupied’ environments could facilitate niche expansion for clownfish species (Bruno *et al*. 2003; Bulleri *et al*. 2016). Sea anemones may create favorable microenvironments (Arossa *et al*. 2021) enabling clownfish larvae to settle in otherwise harsh conditions. These areas could foster new environmental adaptations and driving range expansions (Chen *et al*. 2018; Álvarez *et al*. 2020). This host-mediated niche expansion might have facilitated the clownfish clade’s expansion into the West Indian Ocean around 5 million years ago (Litsios *et al*. 2014), with long-established sea anemones (Titus *et al*., 2019) providing an ecological opportunity for adapted clownfish. This hypothesis highlights the interplay between biotic interactions and abiotic factors in shaping species distributions and evolution. It also suggests sea anemones act as niche constructors, modifying environments for their symbionts, similar to mycorrhizal fungi altering soil for plants or corals building reef habitats.

Consistent with previous research on mutualistic networks (Bascompte & Jordano, 2007), generalist clownfish species showed higher niche overlap with host anemones compared to specialists. This suggests generalist niches are less constrained by biotic interactions, while specialist niches are strongly shaped by the specificity of their mutualistic relationship (Fig. 3). Specialist exhibited larger proportions of unavailable niches, likely due to their tight association with fewer hosts. This makes niche estimations incorporating biotic interactions more likely to differ from environmental-only models (Bateman *et al*. 2012), particularly for specialists linked to rare or patchily distributed hosts. Specialist niche models without accounting for biotic constrains may be prone to overfitting, potentially leading to inaccurate predictions, oversimplified understanding of complex ecological dynamics and misguided conservation efforts (Dormann *et al*. 2018). This is particularly crucial given as specialist species are often more vulnerable to climate change and extinction risks. Conversely, areas of overlapping predicted niches for clownfish and host anemones enhance model robustness, increasing confidence in distribution estimates and validating our understanding of their mutualism and ecological requirements.

While our niche estimations are considered reliable, potential biases stemming from imbalance occurrence data or incorrect biotic associations may exist. UUU parameters can identify areas needing better sampling or reveal misidentified associations, as seen in recent observations of *A. latezonatus, A. chagosensis* and *A. sebae* (Gaboriau *et al*. 2024). These species showed associations with more sea anemone species than previously known, potentially explaining high levels of unavailable niche proportions. *A. sebae*’s distribution, spanning from the Central Indian Ocean to the Eastern Coral Triangle, shows decreasing Unavailability proportions from west to east (100% in Central Indian Ocean Islands, 52% in Andaman, and 14% in Western Coral Triangle). This gradient suggests its western range could be facilitated by novel ecological interactions, indicating a shift in ecological structuring based on different host associations across its distribution. Similar patterns at smaller scales are observed for *A. chagosesis* (40% Unavailability in Central Indian Ocean Islands) and *A. latezonatus* (14% in East Central Australian Shelf and 100% in Lord Howe Island). These analyses provide a valuable tool for conservation strategies by identifying areas or populations requiring increased monitoring efforts.

### Effect of mutualistic interactions on species niche overlaps

Our models provide insights into clownfish interspecific competition dynamics. Competition relies on physical interactions between species, making niche and distribution inferences crucial (Godsoe *et al*. 2015). Our approach enhances clownfish distribution estimations by considering their nested presence within hosts and allows studying interactions while accounting for resource partitioning due to varying host use.

Ecological niche overlaps were high, regardless of mutualistic associations and host use. As such, two clownfish in the same region would have similar environmental suitability despite inhabiting different sea anemones. Tropical reef fishes have narrow environmental niches, with limited tolerance to variations in temperature, salinity, pH, and oxygen levels (Brandl *et al*. 2020; Munday *et al*. 2008). Many species show reduced fitness when conditions deviate from their optimal range (Johansen *et al*. 2014; Rummer *et al*. 2013). Damselfishes exhibit decreased maximum oxygen uptake and aerobic scope at temperatures above 31°C (Habary *et al*. 2017). This specialization makes reef fish sensitive to environmental changes and vulnerable to climate impacts (Munday *et al*. 2008; Brandl *et al*. 2020). Resource-use overlap among clownfish species varied based on shared host anemones and mutualistic behaviours. Species sharing multiple host anemones exhibited greater overlap compared to those sharing fewer hosts. Generalist clownfish, associating with a wider range of anemone hosts, demonstrated higher resource-use overlap than specialists, restricted to fewer host species.

Our analyses revealed an inverse relationship between resource-use overlap and ecological niche overlap among clownfishes. While sea anemone associations are critical for clownfish survival and reproduction (Lubbock 1980; Fautin 1991), factors driving species-specific host preferences remain unclear. We suggest that current mosaic of host preferences arose from long-term competitive dynamics aimed at minimizing resource-use overlaps, .facilitating species coexistence in optimal environments with high ecological niche overlap. This competition avoidance through resource partitioning is observed in various mutualistic systems, such as yucca-yucca moths, acacia-ant, plant-pollinator, and ant-myrmecophyte mutualisms (e.g., Addicott 1998; Palmer *et al*. 2003; Lee *et al*. 2010; Jeavons *et al*. 2020). In clownfishes, host resource-use overlap likely drives interspecific competition, as these species share fundamental ecological characteristics such as trophic position, reproductive behavior, phenology, and social structure. Host use and mutualistic behavior are primary differentiating factors, influencing coloration patterns and species recognition (Gaboriau *et al*. 2024). This suggests host use is a strong predictor of potential competitive dynamics among clownfish species.

While other ecological factors may influence local competition dynamics and the scale and resolution of the niche estimations could limit detection of biotic interactions (e.g. Pearson & Dawson 2003, Araujo & Rozenfeld 2014 and Fontoura *et al*. 2020), our analysis reveals potential effects of mutualistic interactions on clownfish coexistence and competitive dynamics at the regional scale. Our study reveals that generalist-specialist interactions are the most prevalent, with 67 unique interactions, particularly in species-rich environments. These interactions exhibit the lowest resource-use overlap, suggesting they are favoured in saturated communities, reducing competition and enabling species coexistence. Such interactions align with observations in plant-animal mutualistic systems, where generalist-specialist interactions contribute to biodiversity maintenance (Bascompte & Jordano 2007).

### Geographical patterns of resource-use overlap

We observed distinct coexistence patterns among clownfish species across regions (Fig. S16 and Table S5, each linked to contrasting levels of ecological niche and resource-use overlap. These patterns reflect the complex interplay between host specialization and environmental adaptation in shaping distributions, aligning with the observed clownfish evolutionary dynamics (Litsios *et al*. 2014), suggesting a trade-off between host and environmental specialization.

Our findings align with Camp *et al*. (2016), where approximately 25% of clownfishes inhabiting the Coral Triangle were found in interspecific cohabiting groups, where clownfish species richness often exceeded host availability. This region characterized by high species cooccurrence and significant differences between ecological and resource-use overlap, exhibits a common coexistence pattern involving a generalist and a specialist with low resource-use overlap, indicating opportunities for coexistence through niche partitioning (Polechová & Storch 2019). However, we identified two alternative coexistence scenarios in which resource-use overlap was high, suggesting on-going competition.

First, regions at distribution edges such as the Western Pacific, Somali/Arabian Peninsula, and Central and Western Indian Ocean, exhibited high resource-use overlap driven mainly by closely related species. These regions, characterized by more recent colonization events (Litsios *et al*. 2014), likely represent areas of ongoing adaptation to new environmental conditions. These species pairs share host anemones while adapting to different environmental conditions, indicating recent divergence and specialization in distinct ecological niches. Some of those interactions have been reported as never coexisting within the same location, suggesting fine-scale competitive exclusion not captured by our regional-scale analyses.

Second, we found distantly related species, like *A. frenatus* and *A. biaculeatus* or *A. chrysopterus* and *A. percula*, converging at distribution edges and sharing hosts despite evolving in different ecological niches, indicating secondary contact and convergence on common hosts (Gaboriau *et al*. 2024). In these cases, we observed divergence between species in other traits such as coloration, morphology, or positioning relative to the host anemone. This observation aligns with findings by Camp *et al*. (2016). Such trait divergence likely serves to reduce competition when species share host anemones, enabling cohabitation on the same host while maintaining species recognition to prevent hybridization.

Our study reveals complex resource-use overlap dynamics influenced by species richness, ecological overlap, and specialization levels. Generalist-generalist interactions show increased resource-use overlap as ecological overlap and generalist species number rise. Coexistence is most likely in communities with few generalists and moderate ecological overlap. However, generalists’ flexibility in host switching may mitigate competition locally. Specialist-specialist interactions exhibit a different pattern, where a larger number of specialists allows reduced competition if ecological overlap is low. Specialist-generalist interactions strike an optimal balance, supporting high species richness and ecological overlap while maintaining low resource-use overlap. This balance likely promotes species coexistence and may explain the remarkable clownfish diversity in the Coral Triangle, the most environmentally competitive area. These findings underscore the importance of considering specialization levels and ecological factors in understanding biodiversity maintenance mechanisms in complex ecosystems like coral reefs.

## Conclusion

Our study introduces a novel framework for assessing species interactions’ impact on niche estimation, with significant implications for conservation strategies and understanding mutualistic networks. Applied to clownfishes, we revealed the crucial role of resource partitioning in regulating competition, illuminating the evolution of diverse clownfish-sea anemone associations. Our findings support established hypotheses in mutualistic systems, demonstrating how diverse strategies promote ecosystem sustainability and mitigate negative effects in saturated communities. Competition avoidance through resource partitioning emerges as a central mechanism shaping clownfish communities across the Indo-Pacific, aligning with broader ecological principles. While our study highlights importance of resource partitioning, it also raises questions about its evolutionary origins versus dynamic responses to current host composition. This framework advances clownfish ecology understanding and provides a valuable tool for investigating mutualistic interactions broadly. It can be extended to account for various interaction types, contributing to a more comprehensive understanding of biodiversity maintenance in complex ecosystems.

## Supplementary information

## Acknowledgements

We thank Daniele Silvestro, Pablo Duchen and Thibault Latrille for their contribution to the discussions of this study.

## Data availability

All data sets used and produced, figures and R scripts can be retrieved from the DRYAD repository (https://doi.org/10.5061/dryad.2bvq83bv8).

## Code availability

The scripts used for the analyses will be deposited on Dryad upon acceptance of the manuscript, ensuring the reproducibility and accessibility of our research findings.

## Author contribution

AGJ, TG and NS designed the study, AGJ collected the data, developed the implementation, performed the modelling, carried out the analyses, and wrote the initial manuscript. TG and NS contributed to the development of the implementation, the interpretation of the analyses and the writing of the manuscript, with help of all authors.

