## Supplementary Material for "Integrating Biotic Interactions In Niche Analyses Unravels Patterns Of Community Composition in Clownfishes"

[1*]Department of Computational Biology, University of Lausanne, Lausanne, Switzerland.
[2]Department of Ecology and Evolution,University of Lausanne, Lausanne, Switzerland.
[3]Institute of Earth Science Dynamics, University of Lausanne, Lausanne, Switzerland.
[*] indicates co-last authorship.

Contributing authors:;

#### Abstract

Biotic interactions shape the ecology of species and communities, yet their integration into ecological niche modeling methods remains challenging. Despite being a central topic of research for the past decade, the impact of biotic interactions on species distributions and community composition is often overlooked. Mutualistic systems offer ideal case studies for examining the effects of biotic interactions on species niches and community dynamics. This study presents a novel approach to incorporating mutualistic interactions into niche modeling, using the clownfish-sea anemone system. By adapting existing niche quantification frameworks, we developed a method to estimate the partial effects of known interactions and refine ecological niche estimates. This approach allows for a more comprehensive understanding of how mutualistic relationships influence species distributions and community assembly patterns. We also used mutualistic information to investigate the resource-use overlap, identityng patterns of competition within clownfish communities. Our results reveal significant deviations in niche estimates when biotic interactions are considered, particularly for specialist species. Host partitioning among clownfish species reduces resource-use overlap, facilitating coexistence in species-rich habitats and highlighting mutualism's role in promoting and maintaining diversity. We uncover complex dynamics in resource-use overlap among clownfish species, influenced by factors such as species richness, ecological niche overlap, and host specialization. Specialist-generalist interactions strike an optimal balance, supporting high species richness while minimizing competition. These insights enhance our understanding of clownfish biodiversity patterns, demonstrating how diverse mutualistic strategies contribute to diversity build-up and mitigate competitive exclusion in saturated communities. The analytical framework presented has broad applications beyond the clownfish-sea anemone system, potentially extending to a broader range of interactions. It enables a more comprehensive understanding of biodiversity maintenance in complex ecosystems and constitutes a valuable tool for conservation planning and ecosystem management.

**Keywords:** biotic interactions, clownfish, community composition, competition, mutualism, niche, species distribution, spatial ecology

#### Supplementary tables

**Tab. S1**: Spatial Autocorrelation Analysis: Moran's I Statistics and Correlogram Results for Resource-Use Overlap to assess the spatial dependency of resource-use overlap in clownfish-anemone interactions. The analysis is conducted across 20 distance classes (dist.class, measured in km). For each distance class, the table reports the Moran's I coefficient (coef), its associated p-value, and the number of pair comparisons (n). Positive coefficients indicate positive spatial autocorrelation, while negative coefficients suggest negative spatial autocorrelation. The p-values indicate the statistical significance of the observed spatial patterns at each distance class. This analysis helps to quantify the scale and strength of spatial dependencies in the ecological relationship between clownfish and sea anemones.

| dist.class | coef | p.value | n |
|---|---|---|---|
| 3.959 | 0.638 | 0.00E+00 | 1338576 |
| 11.712 | 0.383 | 0.00E+00 | 2080136 |
| 19.465 | 0.12 | 0.00E+00 | 2130268 |
| 27.218 | -0.088 | 1.00E+00 | 1904600 |
| 34.971 | -0.124 | 1.00E+00 | 1521748 |
| 42.724 | -0.087 | 1.00E+00 | 1249776 |
| 50.477 | -0.036 | 1.00E+00 | 826122 |
| 58.23 | -0.007 | 9.97E-01 | 700014 |
| 65.983 | -0.082 | 1.00E+00 | 620488 |
| 73.736 | -0.147 | 1.00E+00 | 450760 |
| 81.489 | -0.092 | 1.00E+00 | 389964 |
| 89.242 | -0.004 | 7.70E-01 | 304982 |
| 96.995 | 0.097 | 1.50E-81 | 255992 |
| 104.748 | 0.157 | 1.13E-137 | 154706 |
| 112.501 | 0.116 | 1.01E-37 | 93076 |
| 120.254 | 0.116 | 3.57E-38 | 42100 |
| 128.007 | -0.118 | 1.00E+00 | 19670 |
| 135.76 | -0.172 | 1.00E+00 | 20220 |
| 143.513 | -0.243 | 1.00E+00 | 7098 |
| 151.266 | -1.037 | 1.00E+00 | 996 |

**Tab. S2**: Results of Cramer's V tests, which assess the statistical significance of changes in species' niche responses when mutualistic interactions are incorporated into the models. For each species, the test compares the original ecological niche to the mutualism-refined niche, quantifying the impact of host availability on niche estimation. Tests were performed across each geographic regions, providing insights into the spatial patterns of niche change due to their mutualistic interactions.

| Species | Province | Cramer statistic | p-value | Species | Province | Cramer statistic | p-value |
|---|---|---|---|---|---|---|---|
| *A. akallopisos* | Andaman | 6.139 | 9.99E-04 | *A. melanopus* | East Central Australian Shelf | 0.199 | 6.81E-01 |
| *A. akallopisos* | Sunda Shelf | 1.883 | 9.49E-02 | *A. melanopus* | Eastern Coral Triangle | 7.578 | 9.99E-04 |
| *A. akallopisos* | Western Indian Ocean | 1.736 | 3.10E-02 | *A. melanopus* | Marshall Gilbert and Ellis Islands | 9.743 | 0.00E+00 |
| *A. akindynos* | East Central Australian Shelf | 7.156 | 2.00E-03 | *A. melanopus* | Northeast Australian Shelf | 1.498 | 1.94E-01 |
| *A. akindynos* | Northeast Australian Shelf | 0.652 | 5.55E-01 | *A. melanopus* | Northwest Australian Shelf | 1.345 | 3.04E-01 |
| *A. akindynos* | Sahul Shelf | 2.531 | 1.10E-02 | *A. melanopus* | Sahul Shelf | 2.662 | 2.20E-02 |
| *A. akindynos* | Tropical Southwestern Pacific | 0.741 | 3.74E-01 | *A. melanopus* | Tropical Northwestern Pacific | 3.239 | 1.50E-02 |
| *A. allardi* | Western Indian Ocean | 0.907 | 1.47E-01 | *A. melanopus* | Tropical Southwestern Pacific | 2.711 | 1.90E-02 |
| *A. barberi* | Tropical Southwestern Pacific | 10.066 | 0.00E+00 | *A. melanopus* | Western Coral Triangle | 4.07 | 9.99E-04 |
| *A. bicinctus* | Central Indian Ocean Islands | 7.563 | 0.00E+00 | *A. nigripes* | Central Indian Ocean Islands | 6.094 | 0.00E+00 |
| *A. bicinctus* | Red Sea and Gulf of Aden | 1.105 | 2.33E-01 | *A. nigripes* | West and South Indian Shelf | 32.128 | 0.00E+00 |
| *A. chagosensis* | Central Indian Ocean Islands | 18.784 | 0.00E+00 | *A. ocellaris* | Andaman | 4.292 | 1.30E-02 |
| *A. chrysogaster* | Western Indian Ocean | 1.205 | 1.69E-01 | *A. ocellaris* | Java Transitional | 5.668 | 5.00E-03 |
| *A. chrysopterus* | Central Polynesia | 4.696 | 7.99E-03 | *A. ocellaris* | Northwest Australian Shelf | 17.409 | 0.00E+00 |
| *A. chrysopterus* | Eastern Coral Triangle | 9.456 | 2.00E-03 | *A. ocellaris* | Sahul Shelf | 7.737 | 0.00E+00 |
| *A. chrysopterus* | Marshall Gilbert and Ellis Islands | 0.834 | 4.19E-01 | *A. ocellaris* | South China Sea | 4.634 | 3.00E-03 |
| *A. chrysopterus* | Northeast Australian Shelf | 0.356 | 8.91E-01 | *A. ocellaris* | South Kuroshio | 8.137 | 9.99E-04 |
| *A. chrysopterus* | Southeast Polynesia | 7.296 | 0.00E+00 | *A. ocellaris* | Sunda Shelf | 0.289 | 5.08E-01 |
| *A. chrysopterus* | Tropical Northwestern Pacific | 0.498 | 6.45E-01 | *A. ocellaris* | Western Coral Triangle | 0.816 | 2.12E-01 |
| *A. chrysopterus* | Tropical Southwestern Pacific | 0.634 | 5.10E-01 | *A. omanensis* | Somali Arabian | 1.356 | 7.29E-02 |
| *A. clarkii* | Andaman | 2.126 | 1.36E-01 | *A. percula* | Eastern Coral Triangle | 1.404 | 1.11E-01 |
| *A. clarkii* | Central Indian Ocean Islands | 0.841 | 3.18E-01 | *A. percula* | Northeast Australian Shelf | 1.981 | 9.99E-02 |
| *A. clarkii* | Eastern Coral Triangle | 1.54 | 1.55E-01 | *A. percula* | Southeast Polynesia | 35.481 | 0.00E+00 |
| *A. clarkii* | Java Transitional | 3.166 | 2.70E-02 | *A. percula* | Tropical Southwestern Pacific | 0.31 | 7.34E-01 |
| *A. clarkii* | Northeast Australian Shelf | 0.56 | 4.83E-01 | *A. perideraion* | Eastern Coral Triangle | 6.971 | 9.99E-04 |
| *A. clarkii* | Northwest Australian Shelf | 0.908 | 3.55E-01 | *A. perideraion* | Java Transitional | 7.53 | 9.99E-04 |
| *A. clarkii* | Sahul Shelf | 4.274 | 1.40E-02 | *A. perideraion* | Northeast Australian Shelf | 1.65 | 1.55E-01 |
| *A. clarkii* | Somali Arabian | 3.792 | 7.99E-03 | *A. perideraion* | Northwest Australian Shelf | 0.914 | 3.65E-01 |
| *A. clarkii* | South China Sea | 1.316 | 6.89E-02 | *A. perideraion* | Sahul Shelf | 0.085 | 8.70E-01 |
| *A. clarkii* | South Kuroshio | 1.575 | 2.72E-01 | *A. perideraion* | South China Sea | 6.873 | 0.00E+00 |
| *A. clarkii* | Sunda Shelf | 0.025 | 9.99E-01 | *A. perideraion* | South Kuroshio | 1.774 | 1.15E-01 |
| *A. clarkii* | Tropical Northwestern Pacific | 2.76 | 4.50E-02 | *A. perideraion* | Sunda Shelf | 1.177 | 1.61E-01 |
| *A. clarkii* | Tropical Southwestern Pacific | 0.559 | 5.69E-01 | *A. perideraion* | Tropical Northwestern Pacific | 0.493 | 4.87E-01 |
| *A. clarkii* | Warm Temperate Northwest Pacific | 2.113 | 6.29E-02 | *A. perideraion* | Tropical Southwestern Pacific | 4.589 | 3.00E-03 |
| *A. clarkii* | West Central Australian Shelf | 13.589 | 0.00E+00 | *A. perideraion* | Western Coral Triangle | 1.903 | 6.19E-02 |
| *A. clarkii* | Western Coral Triangle | 0.415 | 4.60E-01 | *A. polymnus* | Eastern Coral Triangle | 9.192 | 0.00E+00 |
| *A. clarkii* | Western Indian Ocean | 2.906 | 4.70E-02 | *A. polymnus* | Sunda Shelf | 2.699 | 3.80E-02 |
| *A. clarkii* | West and South Indian Shelf | 85.551 | 0.00E+00 | *A. polymnus* | Western Coral Triangle | 0.557 | 3.31E-01 |
| *A. ephippium* | Andaman | 8.134 | 0.00E+00 | *A. polymnus* | South China Sea | 21.869 | 0.00E+00 |
| *A. ephippium* | Java Transitional | 19.274 | 0.00E+00 | *A. rubrocinctus* | Northwest Australian Shelf | 2.591 | 3.50E-02 |
| *A. frenatus* | Andaman | 18.312 | 0.00E+00 | *A. rubrocinctus* | Sahul Shelf | 0.965 | 1.10E-01 |
| *A. frenatus* | Java Transitional | 6.443 | 0.00E+00 | *A. rubrocinctus* | Tropical Southwestern Pacific | 8.953 | 0.00E+00 |
| *A. frenatus* | Northwest Australian Shelf | 11.584 | 0.00E+00 | *A. sandaracinos* | Eastern Coral Triangle | 5.869 | 2.00E-03 |
| *A. frenatus* | South Kuroshio | 10.331 | 0.00E+00 | *A. sandaracinos* | Sunda Shelf | 0.087 | 8.39E-01 |
| *A. frenatus* | Sunda Shelf | 0.809 | 1.06E-01 | *A. sandaracinos* | Western Coral Triangle | 3.171 | 1.20E-02 |
| *A. frenatus* | Warm Temperate Northwest Pacific | 22.459 | 0.00E+00 | *A. sebae* | Andaman | 7.131 | 2.00E-03 |
| *A. frenatus* | Western Coral Triangle | 2.955 | 2.00E-03 | *A. sebae* | Western Coral Triangle | 1.646 | 5.59E-02 |
| *A. fuscocaudatus* | Western Indian Ocean | 2.74 | 7.99E-03 | *A. sebae* | Central Indian Ocean Islands | 30.086 | 0.00E+00 |
| *A. latezonatus* | East Central Australian Shelf | 9.776 | 0.00E+00 | *A. tricinctus* | Tropical Southwestern Pacific | 1.705 | 1.03E-01 |
| *A. latezonatus* | Lord Howe and Norfolk Islands | 82.818 | 0.00E+00 | *A. biaculeatus* | Eastern Coral Triangle | 7.085 | 9.99E-04 |
| *A. latifasciatus* | Western Indian Ocean | 9.371 | 0.00E+00 | *A. biaculeatus* | Java Transitional | 16.195 | 0.00E+00 |
| *A. leucokranos* | Eastern Coral Triangle | 14.643 | 0.00E+00 | *A. biaculeatus* | Northeast Australian Shelf | 4.628 | 0.00E+00 |
| *A. mccullochi* | Lord Howe and Norfolk Islands | 18.319 | 0.00E+00 | *A. biaculeatus* | Sahul Shelf | 7.203 | 9.99E-04 |
| | | | | *A. biaculeatus* | Sunda Shelf | 4.038 | 4.00E-03 |
| | | | | *A. biaculeatus* | Western Coral Triangle | 2.204 | 1.40E-02 |

**Tab. S3**: Comparison of niche metrics between generalist and specialist clownfish species. The table presents median (Q2) and interquartile range (IQR) values for various niche and distribution metrics. Statistical comparisons between generalists and specialists are shown with test statistics and *p*-values. Metrics include proportions of unavailable, unoccupied, and used niche/distribution space, as well as measures of niche shift (Centroid Shift, Environmental Shift) and niche characteristics (Niche Dissimilarity, Niche Breadth difference) when accounting for mutualistic interactions.

| | Generalist | | Specialist | | | |
|---|---|---|---|---|---|---|
| | Q2 | IQR | Q2 | IQR | statistic | *p*-value |
| Unavailable niche | 0.0227 | 0.0779 | 0.1828 | 0.3378 | 23.0612 | 1.57E-06 |
| Unoccupied niche | 0.1492 | 0.1787 | 0.0866 | 0.239 | 2.1735 | 1.40E-01 |
| Used niche | 0.8071 | 0.1644 | 0.629 | 0.3338 | 17.1291 | 3.49E-05 |
| Unavailable distribution | 0.0009 | 0.0136 | 0.0414 | 0.1974 | 18.5174 | 1.68E-05 |
| Unoccupied distribution | 0.0114 | 0.0365 | 0.0213 | 0.0467 | 0.0138 | 9.07E-01 |
| Used distribution | 0.9728 | 0.0624 | 0.8775 | 0.209 | 19.6073 | 9.51E-06 |
| Centroid Shift | 0.0658 | 0.0922 | 0.1423 | 0.1877 | 20.681 | 5.43E-06 |
| Environmental Shift | 0.0102 | 0.0141 | 0.025 | 0.0475 | 17.7037 | 2.58E-05 |
| Niche Dissimilarity | 0.8395 | 0.0985 | 0.7265 | 0.1458 | 39.9866 | 2.56E-10 |
| Niche Breadth difference | 0.0137 | 0.0534 | 0.0914 | 0.1409 | 14.1985 | 1.65E-04 |

**Tab. S4**: Spatial Autoregressive Models Examining Resource-Use Overlap Across Species Interaction Types. Results of spatial autoregressive models analyzing the relationship between resource-use overlap, number of species, and ecological overlap for different types of species interactions. Models are shown for all interactions combined, generalist-generalist interactions, generalist-specialist interactions, and specialist-specialist interactions. Fixed effects estimates (Est), standard errors (SE), and t-values are provided, along with random effects parameters (nu, rho, lambda), residual variance (phi), and model fit statistics (log-likelihood). The spatial autoregressive components account for potential spatial autocorrelation in the data.

| **Fixed effects** | **All interactions** | | | **Generalist Generalist** | | | **Generalist-Specialist** | | | **Specialist Specialist** | | |
|---|---|---|---|---|---|---|---|---|---|---|---|---|
| | Est | SE | t | Est | SE | t | Est | SE | t | Est | SE | t |
| (Intercept) | 0.017 | 0.028 | 0.606 | 0.03 | 0.026 | 1.156 | 0.076 | 0.047 | 1.607 | 0.017 | 0.023 | 0.742 |
| num. species | -0.013 | 0.001 | -10.428 | 0.013 | 0.002 | 6.189 | -0.011 | 0.001 | -12.482 | -0.009 | 0.001 | -7.175 |
| ecological overlap | 0.668 | 0.011 | 59.096 | 0.415 | 0.008 | 52.324 | 0.406 | 0.008 | 50.325 | 0.701 | 0.015 | 46.463 |
| num. species x ecological overlap | -0.015 | 0.002 | -7.934 | 0 | 0.002 | 0.049 | -0.001 | 0.001 | -0.851 | 0.005 | 0.005 | 1.057 |
| **Random effects** | | | | | | | | | | | | |
| nu | 0.602 | | | 0.697 | | | 0.614 | | | 0.455 | | |
| rho | 0.077 | | | 0.09 | | | 0.038 | | | 0.048 | | |
| lambda (x+y) | 0.009 | | | 0.007 | | | 0.012 | | | 0.003 | | |
| **Residual variance** | | | | | | | | | | | | |
| phi | 1.00E-06 | | | 1.00E-06 | | | 6.00E-06 | | | 1.00E-06 | | |
| **Likelihood values** | | | | | | | | | | | | |
| logLik | 5286.622 | | | 4525.269 | | | 5970.978 | | | 4474.289 | | |

**Tab. S5**: Averaged values of clownfish and sea anemone species richness, along with various estimated clownfish ecological niche and resource-use overlap metrics, aggregated at the marine province level. It is divided into four main sections: All interactions, Generalist-Generalist interactions, Generalist-Specialist interactions, and Specialist-Specialist interactions. Each section presents key metrics characterizing regional patterns of clownfish species richness, number of interspecific interactions, and spatial overlap proportion (the fraction of locations with co-occurring clownfish species), host-sharing proportion (the fraction of overlapping locations where species utilize common host anemones), ecological overlap (similarity in environmental preferences), and resource-use overlap (extent to which clownfish share host resources).

| Interaction Type | Metric | And | CIO | ECA | ECT | JT | MGE | NEA | NWA | SS | SA | SCS | SK | SuS | TNP | TSP | WTN | WCT | WIO |
|---|---|---|---|---|---|---|---|---|---|---|---|---|---|---|---|---|---|---|---|
| All interactions | Clownfish richness | 5.381 | 3.705 | 2.742 | 8.101 | 5.546 | 2.000 | 6.811 | 5.297 | 5.983 | 2.000 | 2.699 | 3.289 | 7.093 | 3.898 | 7.330 | 2.000 | 8.326 | 5.345 |
| | Host richness | 3.489 | 2.826 | 5.689 | 6.997 | 6.677 | 3.950 | 7.883 | 4.593 | 6.136 | 2.913 | 1.965 | 1.906 | 9.676 | 4.884 | 5.068 | 3.758 | 9.145 | 7.316 |
| | Num. generalist | 0.924 | 0.965 | 1.000 | 5.481 | 2.769 | 1.000 | 5.813 | 2.749 | 2.312 | 1.000 | 0.000 | 2.781 | 2.920 | 2.942 | 6.397 | 1.000 | 3.902 | 2.846 |
| | Num. specialists | 4.457 | 2.739 | 1.742 | 2.619 | 2.777 | 1.000 | 0.999 | 2.548 | 3.671 | 1.000 | 2.699 | 0.508 | 4.172 | 0.956 | 0.932 | 1.000 | 4.424 | 2.500 |
| | Spatial overlap | 0.822 | 0.875 | 0.828 | 0.843 | 0.873 | 1.000 | 0.958 | 0.812 | 0.758 | 1.000 | 0.800 | 0.655 | 0.792 | 0.956 | 0.849 | 1.000 | 0.874 | 0.816 |
| | Host-sharing | 0.435 | 0.998 | 1.000 | 0.771 | 0.644 | 1.000 | 0.902 | 0.772 | 0.838 | 1.000 | 1.000 | 0.831 | 0.533 | 1.000 | 0.975 | 1.000 | 0.612 | 0.966 |
| | Ecological overlap | 0.271 | 0.572 | 0.556 | 0.499 | 0.432 | 0.651 | 0.746 | 0.417 | 0.385 | 0.646 | 0.480 | 0.354 | 0.296 | 0.811 | 0.670 | 0.786 | 0.435 | 0.631 |
| | Resource-use overlap | 0.108 | 0.402 | 0.281 | 0.214 | 0.201 | 0.470 | 0.332 | 0.200 | 0.154 | 0.507 | 0.416 | 0.191 | 0.098 | 0.524 | 0.386 | 0.154 | 0.138 | 0.259 |
| Generalist-Generalist | Num. of interactions | 0.000 | 0.000 | 0.000 | 12.978 | 2.550 | 0.000 | 14.304 | 2.605 | 1.942 | 0.000 | 0.000 | 2.609 | 2.858 | 2.884 | 17.678 | 0.000 | 5.736 | 2.742 |
| | Spatial overlap | 0.000 | 0.000 | 0.000 | 0.424 | 0.223 | 0.000 | 0.697 | 0.207 | 0.103 | 0.000 | 0.000 | 0.704 | 0.134 | 0.519 | 0.750 | 0.000 | 0.201 | 0.233 |
| | Host-sharing | 0.000 | 0.000 | 0.000 | 0.499 | 0.328 | 0.000 | 0.771 | 0.271 | 0.126 | 0.000 | 0.000 | 0.823 | 0.256 | 0.519 | 0.760 | 0.000 | 0.316 | 0.249 |
| | Ecological overlap | 0.000 | 0.000 | 0.000 | 0.610 | 0.721 | 0.000 | 0.834 | 0.654 | 0.341 | 0.000 | 0.000 | 0.599 | 0.739 | 0.825 | 0.663 | 0.000 | 0.779 | 0.703 |
| | Resource-use overlap | 0.000 | 0.000 | 0.000 | 0.328 | 0.341 | 0.000 | 0.417 | 0.394 | 0.143 | 0.000 | 0.000 | 0.333 | 0.297 | 0.612 | 0.407 | 0.000 | 0.270 | 0.326 |
| Generalist-Specialist | Num. of interactions | 4.254 | 2.664 | 1.742 | 15.028 | 7.835 | 1.000 | 5.810 | 7.327 | 8.849 | 1.000 | 0.000 | 1.320 | 12.363 | 2.855 | 6.098 | 1.000 | 17.509 | 7.336 |
| | Spatial overlap | 0.338 | 0.539 | 0.753 | 0.507 | 0.580 | 1.000 | 0.303 | 0.630 | 0.521 | 1.000 | 0.000 | 0.296 | 0.567 | 0.481 | 0.250 | 1.000 | 0.557 | 0.616 |
| | Host-sharing | 0.632 | 0.538 | 0.753 | 0.477 | 0.350 | 1.000 | 0.229 | 0.665 | 0.542 | 1.000 | 0.000 | 0.177 | 0.628 | 0.481 | 0.240 | 1.000 | 0.562 | 0.596 |
| | Ecological overlap | 0.550 | 0.640 | 0.613 | 0.460 | 0.230 | 0.651 | 0.526 | 0.426 | 0.386 | 0.646 | 0.000 | 0.110 | 0.346 | 0.797 | 0.691 | 0.786 | 0.437 | 0.604 |
| | Resource-use overlap | 0.177 | 0.326 | 0.311 | 0.145 | 0.054 | 0.470 | 0.118 | 0.152 | 0.130 | 0.507 | 0.000 | 0.049 | 0.079 | 0.435 | 0.324 | 0.154 | 0.097 | 0.182 |
| Specialist-Specialist | Num. of interactions | 8.074 | 2.583 | 0.742 | 2.342 | 2.713 | 0.000 | 0.000 | 2.241 | 5.122 | 0.000 | 2.399 | 0.000 | 6.956 | 0.000 | 0.000 | 0.000 | 8.204 | 2.160 |
| | Spatial overlap | 0.662 | 0.461 | 0.247 | 0.070 | 0.197 | 0.000 | 0.000 | 0.162 | 0.375 | 0.000 | 1.000 | 0.000 | 0.300 | 0.000 | 0.000 | 0.000 | 0.242 | 0.151 |
| | Host-sharing | 0.368 | 0.462 | 0.247 | 0.024 | 0.323 | 0.000 | 0.000 | 0.064 | 0.332 | 0.000 | 1.000 | 0.000 | 0.116 | 0.000 | 0.000 | 0.000 | 0.122 | 0.155 |
| | Ecological overlap | 0.132 | 0.505 | 0.442 | 0.173 | 0.750 | 0.000 | 0.000 | 0.151 | 0.403 | 0.000 | 0.480 | 0.000 | 0.089 | 0.000 | 0.000 | 0.000 | 0.223 | 0.637 |
| | Resource-use overlap | 0.073 | 0.477 | 0.221 | 0.059 | 0.500 | 0.000 | 0.000 | 0.151 | 0.207 | 0.000 | 0.416 | 0.000 | 0.068 | 0.000 | 0.000 | 0.000 | 0.139 | 0.422 |

Province abbreviations: **And** = Andaman, **CIO** = Central Indian Ocean Islands, **ECA** = East Central Australian Shelf, **ECT** = Eastern Coral Triangle, **JT** = Java Transitional, **MGE** = Marshall Gilbert and Ellis Islands, **NEA** = Northeast Australian Shelf, **NWA** = Northwest Australian Shelf, **SS** = Sahul Shelf, **SA** = Somali/Arabian, **SCS** = South China Sea, **SK** = South Kuroshio, **SuS** = Sunda Shelf, **TNP** = Tropical Northwestern Pacific, **TSP** = Tropical Southwestern Pacific, **WTN** = Warm Temperate Northwest Pacific, **WCT** = Western Coral Triangle, **WIO** = Western Indian Ocean.

**Tab. S6**: Description of Models Used in Sensitivity Analysis of Clownfish-Anemone Associations

| **Model Label** | **Description of Clownfish-Anemone Interaction Scenario** |
|---|---|
| I. All | Unrestricted: All clownfish interact with all host anemones (no habitat partitioning) |
| II. Generalist | Generalist behavior: All clownfish interact with more than two host species |
| III. rRrGrN | Fully randomized: Randomized regional and global interactions with random mutualistic behaviors |
| IV. rRrGoN | Regionally constrained random: Randomized regional interactions maintaining interaction numbers, but with random host assignments |
| V. Observed | Empirical: Known and established interactions based on field observations (reference model, highlighted in red) |
| VI. rRoGoN | Globally constrained random: Known global interactions with randomized regional subsets |
| VII. Specialist | Specialist behavior: All clownfish interact with fewer than three host species |
| VIII. One | Extreme specialization: Each clownfish interacts with only one unique host species (maximum habitat partitioning) |

#### Supplementary figures

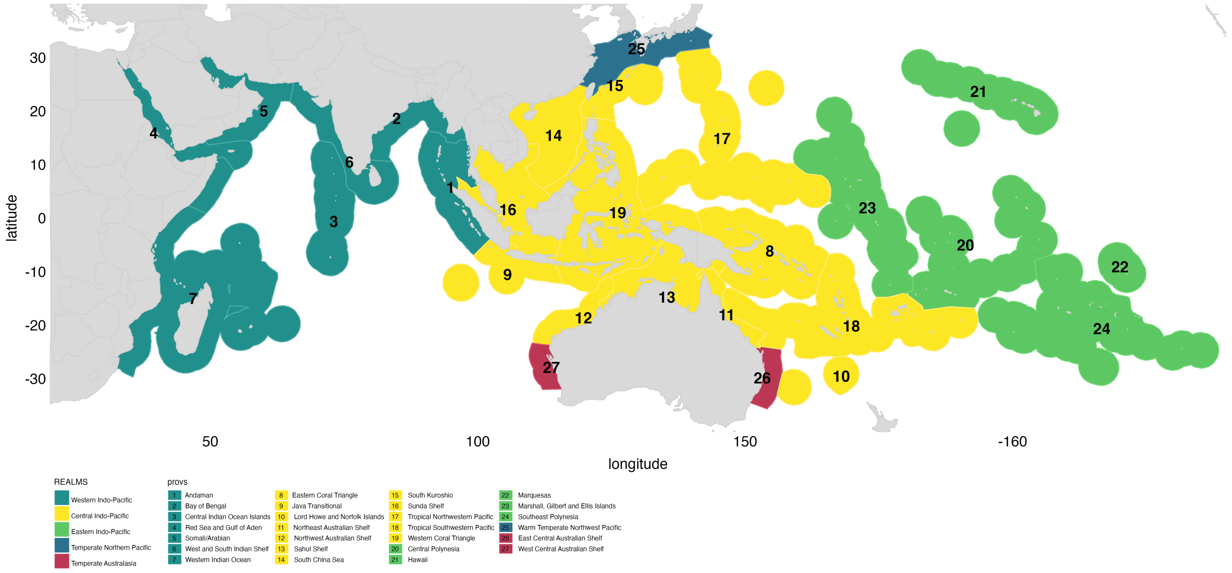


**Fig. S1**: **Global Distribution of Marine Ecoregions of the World (MEOW) Provinces and Realms** Geographic distribution of marine provinces and realms according to the MEOW classification system. Distinct colors represent different marine realms as indicated in the legend (bottom right). Marine provinces, shown as lighter-colored polygons within each realm, are numerically labeled and correspond to the legend (bottom). This visualization provides a comprehensive overview of the biogeographic framework used in analyzing clownfish-anemone interactions across various marine ecosystems.

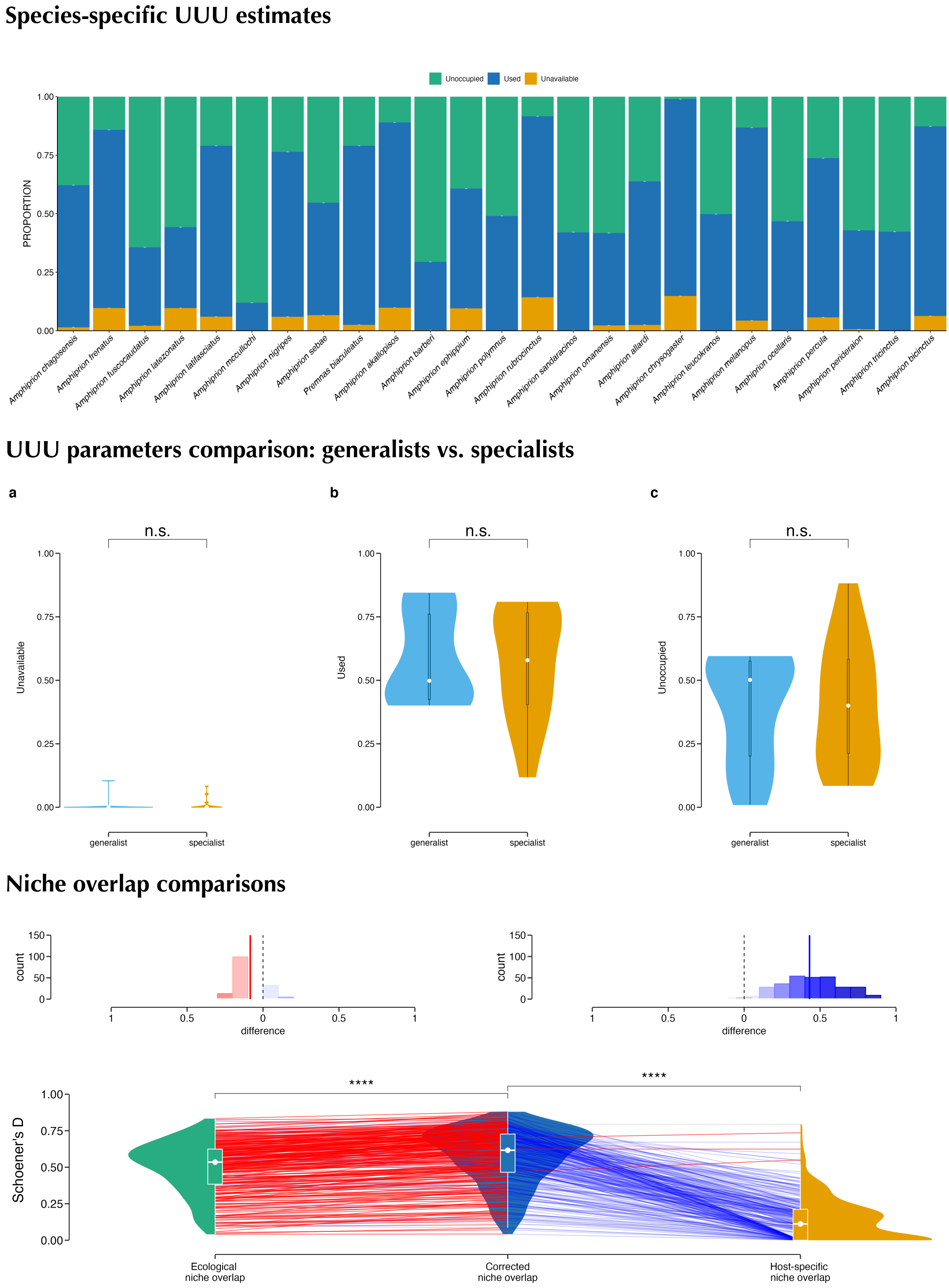


**Fig. S2**: **Global-Scale Analysis of Clownfish-Anemone Symbioses: UUU Estimates and Niche Overlaps** Results from global-scale models of clownfish-anemone symbioses, offering a comparative perspective to the regional-scale analyses presented in the main text. (A) Species-specific estimates of Unavailable, Unoccupied, and Used (UUU) environmental space at a global scale. The lower proportions of Unavailability compared to regional models reflect the coarser resolution of global analyses. (B) Comparison of UUU parameters between generalist and specialist clownfish species at the global scale. Unlike regional models, global analyses show no significant differences between these ecological strategies, highlighting the scale-dependent nature of these patterns. (C) Ecological niche, mutualism-refined niche, and resource-use overlaps among clownfish species at the global scale. These results demonstrate consistency with regional-scale models, supporting the robustness of our findings across different spatial resolutions. The global-scale analysis provides valuable insights into the scale-dependence of clownfish-anemone symbioses. While some patterns, such as niche overlaps, remain consistent across scales, others, like the distinction between generalist and specialist strategies, become less pronounced at the global level.

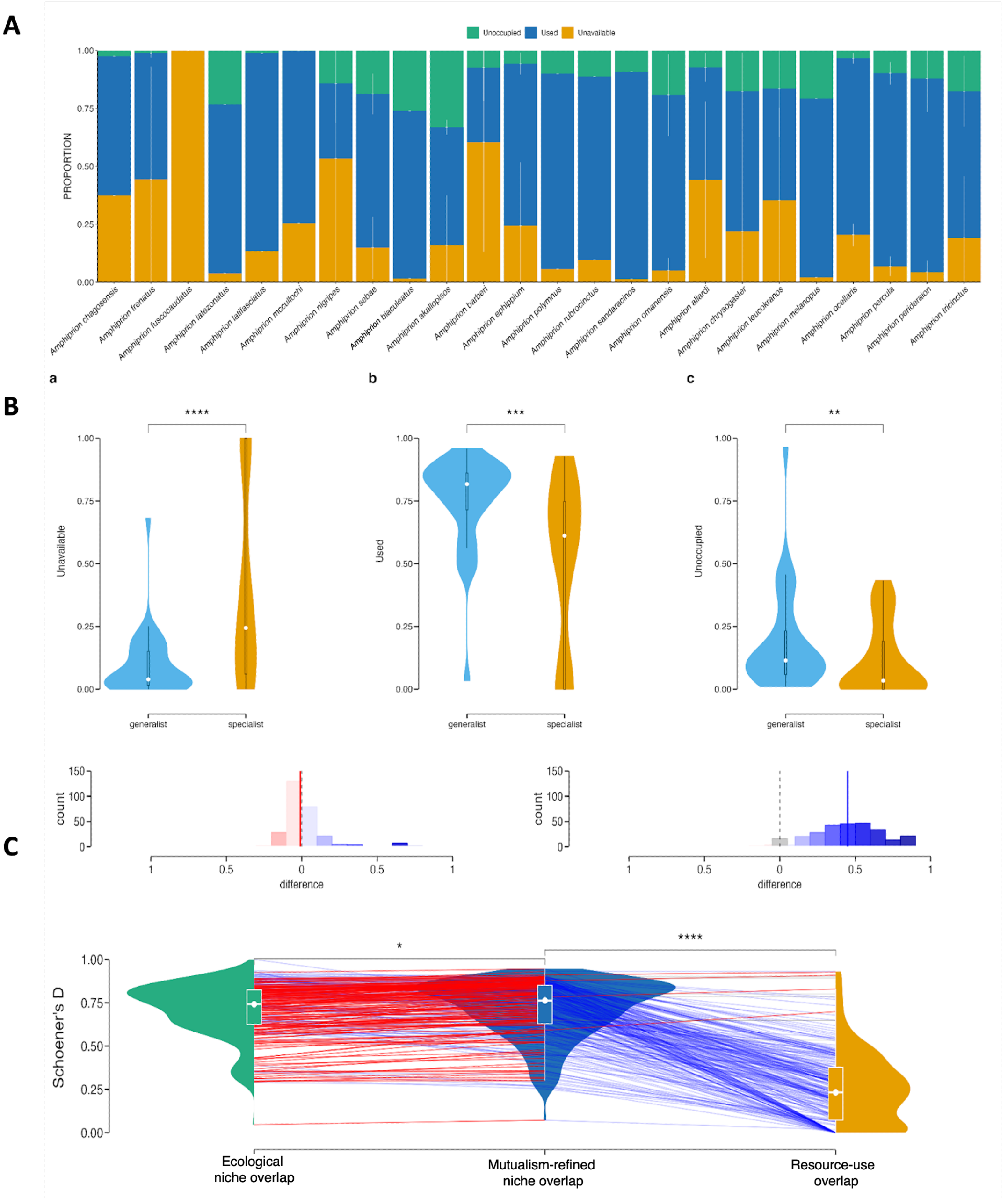


**Fig. S3**: **Robustness of Clownfish-Anemone Symbiosis Models to Environmental Variable Selection** Results of ecological niche models for clownfish-anemone symbioses conducted without prior environmental variable selection, demonstrating the robustness of our findings. The panels correspond to key analyses presented in the main text: a) Comparison to Fig. 2: Niche and spatial availability patterns b) Comparison to Fig. 3: Host utilization and niche overlap relationships c) Comparison to Fig. 4: Biogeographical patterns of symbiotic associations Each panel shows how the results from these comprehensive models, which include all available environmental variables, align with models presented in the main text where variable selection was performed. The consistency between these approaches validates the stability of our findings across different modeling strategies. Observed similarities include patterns in niche and spatial availability, relationships between host utilization and niche overlap, and broad-scale biogeographical trends in symbiotic associations. This analysis underscores the reliability of our conclusions regarding the ecological and evolutionary dynamics of clownfish-anemone mutualisms, demonstrating that they are not artifacts of variable selection procedures but reflect robust underlying patterns in the data.

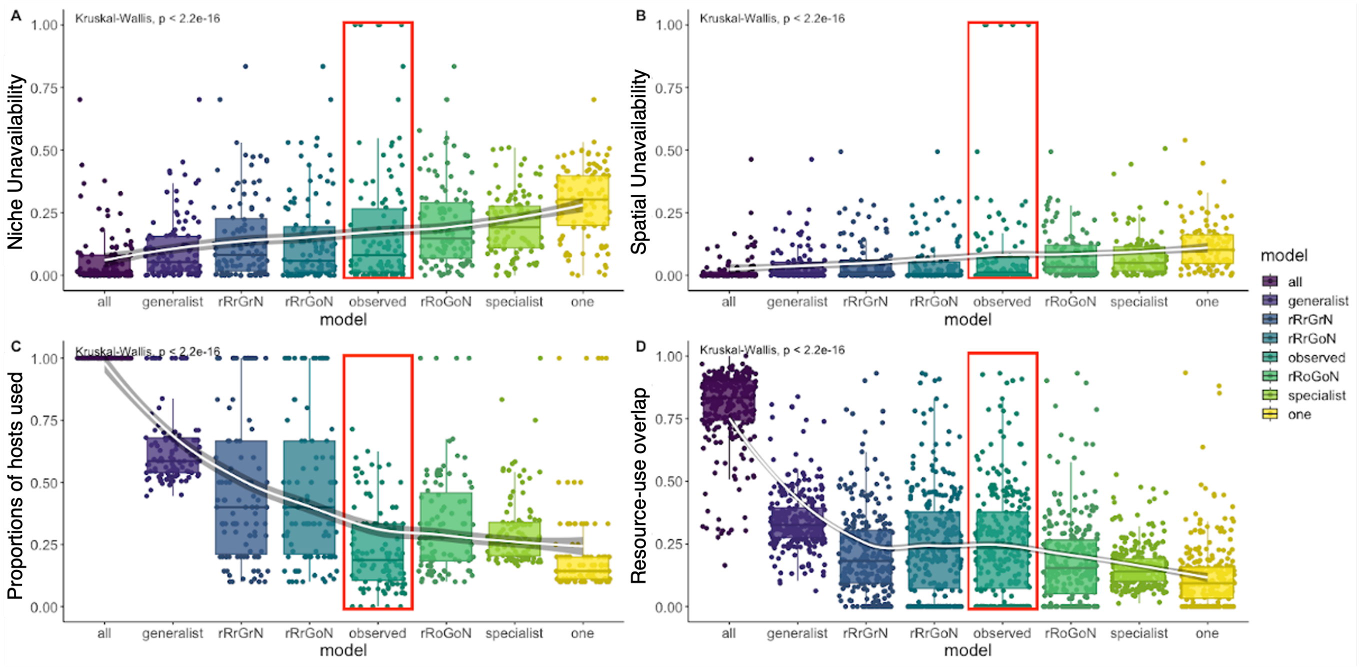


**Fig. S4**: **Sensitivity Analysis of Clownfish-Anemone Association Matrices on Ecological Metrics** Comprehensive sensitivity analysis examining how different clownfish-anemone association matrices affect key ecological metrics. Each panel displays boxplots representing the median and interquartile range (IQR) of a specific metric across different association scenarios: A) Niche unavailability B) Spatial unavailability C) Proportion of hosts used relative to available sea anemones per region D) Resource-use overlap The association matrices, arranged from least to most restrictive habitat partitioning, are: - **All:** Every clownfish interacts with all hosts (no partitioning) - **Generalist:** All species behave as generalists (>2 hosts) - **rRrGrN:** Randomized regional and global interactions with random mutualistic behaviors - **rRrGoN:** Randomized regional interactions maintaining the number of interactions but with random hosts - **Observed:** Known and established interactions (highlighted in red) - **rRoGoN:** Known global interactions with randomized regional interactions - **Specialist:** All species behave as specialists (<3 hosts) - **One:** Each clownfish interacts with only one unique host (maximum partitioning) A Generalized Additive Model (GAM) regression (white line with shaded confidence intervals) illustrates the trend from no habitat partitioning to complete partitioning. This analysis reveals how assumptions about clownfish-anemone associations impact our understanding of niche dynamics, spatial distributions, host utilization, and resource overlap in these symbiotic relationships. The observed interactions (red box) can be compared against various theoretical and randomized scenarios.

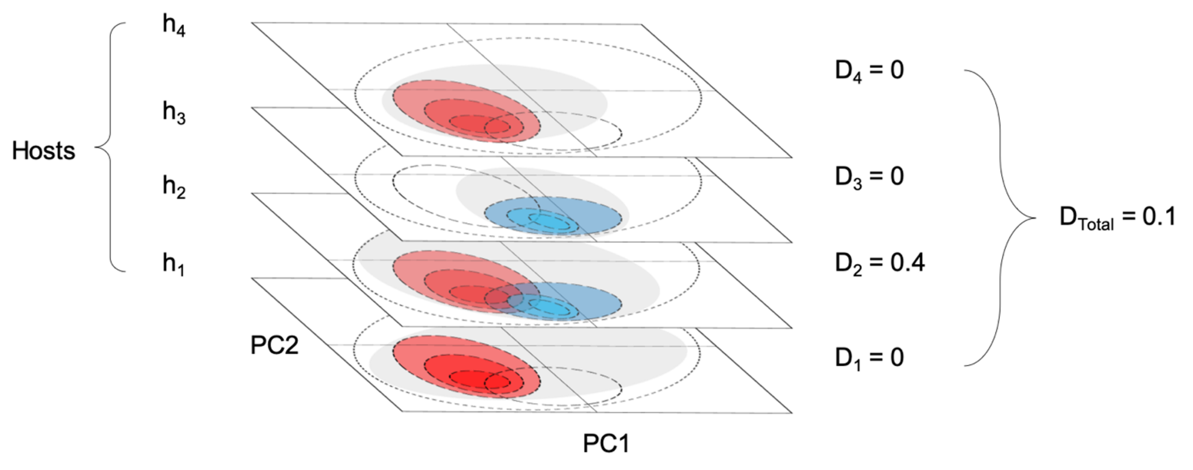


**Fig. S5**: **Conceptual Framework for Estimating Resource-Use Overlap in Clownfish-Anemone Interactions** Multi-step process for calculating resource-use overlap between clownfish species, accounting for their differential host use. First, each clownfish species' environmental niche is projected onto a 2D space derived from PCA, represented by dashed ellipses that are color-coded by species. This projection is then replicated across multiple layers, each representing a host anemone species interacting with either clownfish species. Next, each host anemone's niche is projected onto its respective layer as grey ellipses, while each clownfish species' niche is projected only onto the layers corresponding to its host anemones. The clownfish niche response is weighted by the host niche response in each layer, following equation (1) in the supplementary equations. Schoener's D metric is then used to calculate niche overlap between clownfish species on each layer, and the mean of these layer-specific overlap values represents the overall resource-use overlap. This method captures the complexity of host-specific interactions in determining resource partitioning among clownfish species.

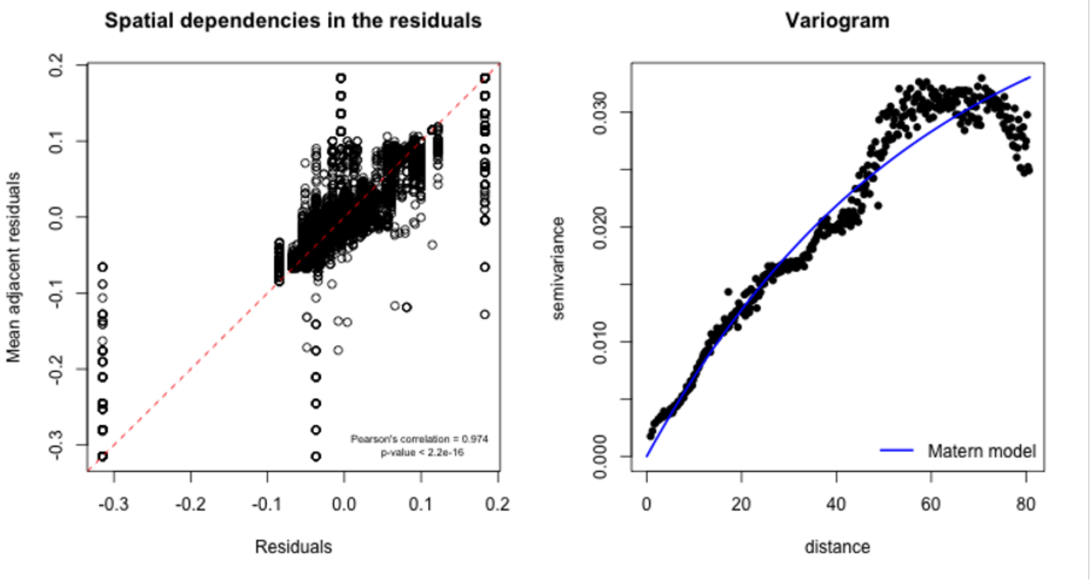


**Fig. S6**: **Spatial Autocorrelation and Variogram Analysis of Clownfish Resource-Use Overlap** The left panel displays a correlation plot between the average values of neighboring locations (grid cells) and the residuals from a generalized linear model (GLM). This analysis serves as a diagnostic tool for detecting spatial autocorrelation in the data. A strong correlation would indicate the presence of spatial dependencies not accounted for by the GLM, justifying the need for spatial modeling techniques. The right panel shows a variogram, which quantifies the variability between data points as a function of distance. The variogram is fitted with a Matérn covariance function, represented by the solid line. This function captures the spatial structure of the data, including the range of spatial dependency and the degree of spatial smoothness. The fitted Matérn function was subsequently used as an input for the spatial generalized linear mixed model (GLMM), ensuring that the final model accurately accounts for the observed spatial patterns in clownfish resource-use overlap.

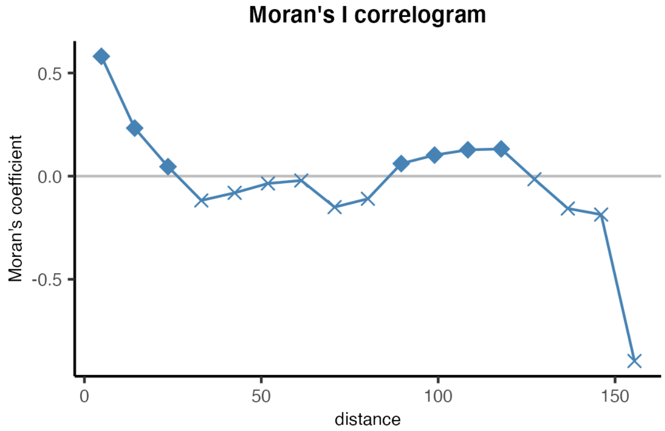


**Fig. S7**: **Spatial Autocorrelation Analysis: Moran's I Correlogram for Clownfish Resource-Use Overlap** This figure presents a Moran's I correlogram for visualizing and quantifying spatial autocorrelation across different distance classes in the clownfish resource-use overlap data. The x-axis represents increasing distance classes, while the y-axis shows the Moran's I coefficient values. Filled diamonds indicate distance classes with statistically significant spatial autocorrelation, whereas crosses denote non-significant distance classes. Positive Moran's I values suggest positive spatial autocorrelation (similar values cluster together), while negative values indicate negative spatial autocorrelation (dissimilar values cluster together). The correlogram's shape reveals the spatial structure of the data, with the decline in Moran's I values over distance potentially indicating the scale at which spatial processes influence clownfish-anemone interactions. This analysis complements the variogram analysis (Figure S10) by providing a clear visualization of how spatial autocorrelation changes with distance.

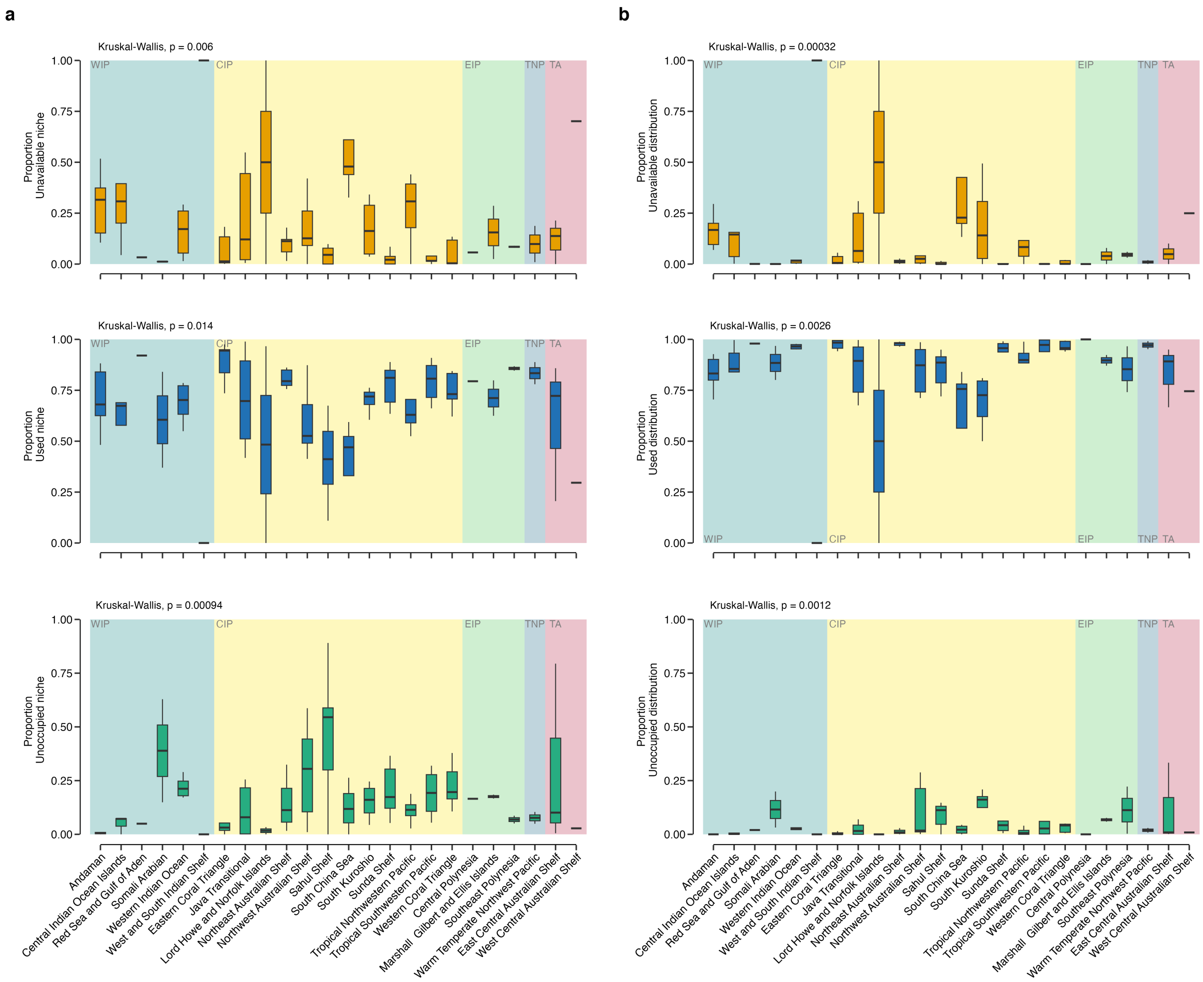


**Fig. S8**: **Distribution of Niche and Spatial UUU Parameters Across Marine Provinces** Proportional distribution of Unavailable, Used, and Unoccupied (UUU) parameters across marine provinces for both niche and spatial dimensions of clownfish-anemone interactions. **a) Niche UUU Proportions:** The top, middle, and bottom panels represent the proportions of Unavailable, Used, and Unoccupied niches, respectively. Each proportion is calculated as the number of environments in a specific UUU category divided by the total number of environments available in that province. **b) Spatial UUU Proportions:** Similarly arranged, these panels show the proportions of Unavailable, Used, and Unoccupied distributions in geographical space. Each proportion is calculated as the number of geographical locations representing environments in a specific UUU category divided by the total number of locations representing all available environments in that province. For each panel, a Kruskal-Wallis test *p*-value is provided in the top-left corner, indicating the statistical significance of differential ecological composition across realms. This quantifies how environmental availability, utilization, and potential for expansion vary across different marine biogeographic regions.

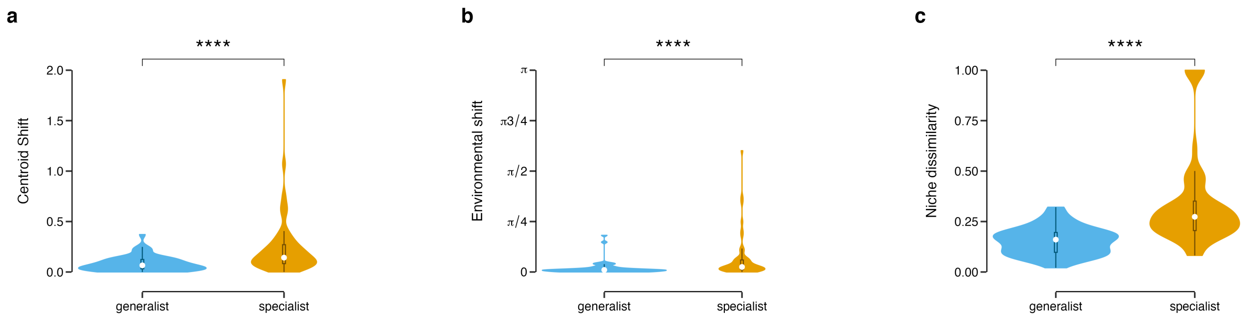


**Fig. S9**: **Impact of Biotic Interactions on Niche Parameters: Generalists vs. Specialists** Effects of incorporating explicit biotic interactions into ecological niche models for generalist and specialist clownfish species. Three key niche parameters are examined: Centroid Shift, which measures the magnitude of change in the central tendency of the species' niche; Environmental Shift, representing the extent of alteration in the overall environmental space occupied by the species; and Niche Dissimilarity, which quantifies the degree of difference between the original and interaction-corrected niches. Violin plots display the distribution of data for each parameter, with wider sections indicating a higher frequency of observations. The comparison between generalists and specialists reveals how species' ecological strategies influence niche adjustments when accounting for biotic interactions. Statistical significance levels are indicated as follows: n.s. (not significant), * ($p < 0.05$), ** ($p < 0.01$), *** ($p < 0.001$), and **** ($p < 0.0001$).

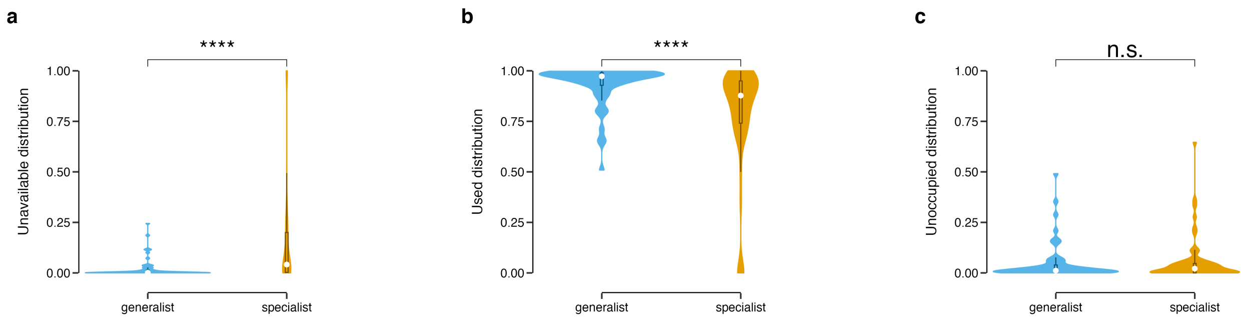


**Fig. S10**: **Comparison of Spatial UUU Parameters Between Generalist and Specialist Clownfish Species** Differences in spatial Unavailable, Used, and Unoccupied (UUU) parameters between generalist and specialist clownfish species. Panel (a) depicts the Unavailable spatial distribution, representing areas that are environmentally suitable but inaccessible. Panel (b) shows the Used spatial distribution, indicating areas that are both suitable and occupied. Panel (c) represents the Unoccupied spatial distribution, highlighting areas that are accessible but environmentally unsuitable. Violin plots are used to visualize the distribution of data for each parameter, with the width of each plot indicating the frequency of observations at different values. The comparison between generalists and specialists provides insights into how ecological specialization influences the spatial patterns of habitat utilization and potential distribution in clownfish-anemone symbioses. Statistical significance of the differences between generalists and specialists is denoted as follows: n.s. (not significant), * ($p < 0.05$), ** ($p < 0.01$), *** ($p < 0.001$), and **** ($p < 0.0001$).

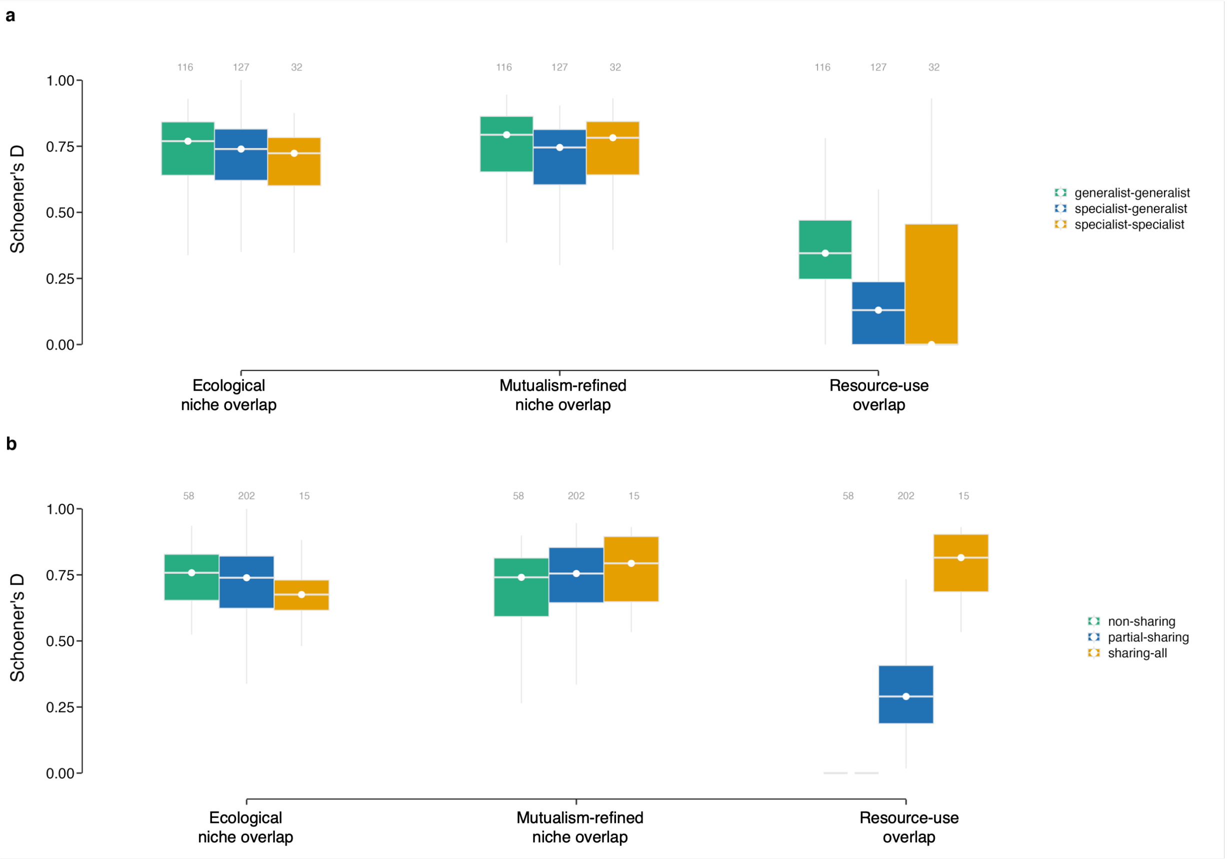


**Fig. S11**: **Influence of Mutualistic Behavior and Host Sharing on Interspecific Niche Overlap in Clownfish Species** Intensity of competition among clownfish species across three niche levels: environmental, mutualism-refined, and resource-use. Panel (a) illustrates the effect of mutualistic behavior on interspecific niche overlap for each niche level. Panel (b) demonstrates how the degree of host anemone sharing (non-sharing, partial-sharing, and sharing-all) influences niche overlap across the three niche levels. Boxplots represent the distribution of pairwise species Schoener's D overlap values per marine province. The colors of the boxes correspond to different interaction types or levels of host sharing, as detailed in the legend on the right. Grey numbers above each boxplot indicate the sample size for each category. This analysis provides insights into how symbiotic relationships and host specificity modulate niche overlap and potential competition among clownfish species. The comparison across niche levels reveals the importance of considering biotic interactions and resource partitioning in understanding the ecological dynamics of clownfishes.

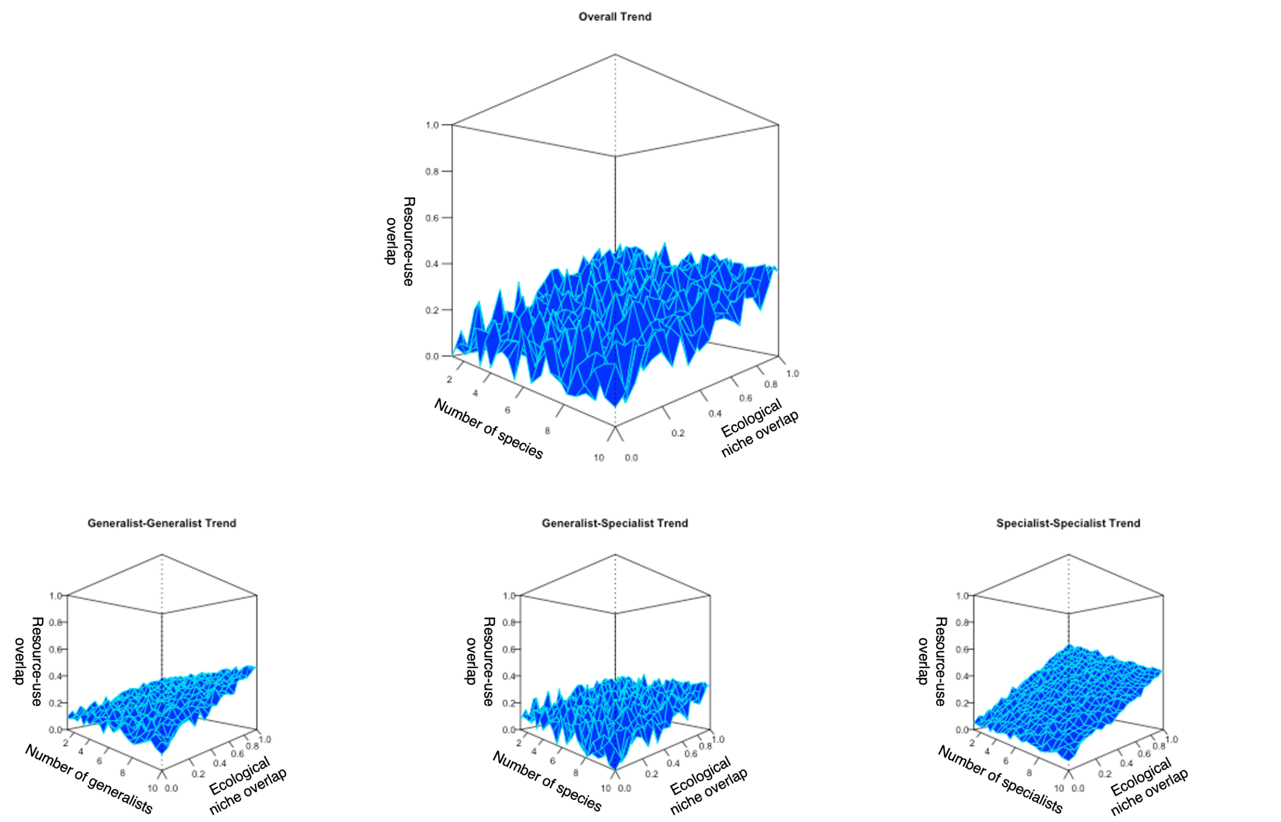


**Fig. S12**: **Spatial Generalized Linear Mixed Models of Resource-Use Overlap in Clownfish Species** Predictions of resource-use overlap as a function of species richness and ecological niche overlap intensity, derived from spatial generalized linear mixed models (GLMMs). The analysis is broken down into four panels: (a) depicts the overall trend across all clownfish species interactions, while panels (b), (c), and (d) show specific trends for Generalist-Generalist, Generalist-Specialist, and Specialist-Specialist interactions, respectively. Each panel illustrates how resource-use overlap (z-axis) varies with the number of species (x-axis) and the intensity of ecological niche overlap (y-axis). These models account for spatial autocorrelation in the data, providing a more accurate representation of the ecological relationships. The comparison across different interaction types reveals how ecological strategies (generalist vs. specialist) influence patterns of resource partitioning and potential competition among clownfish species.

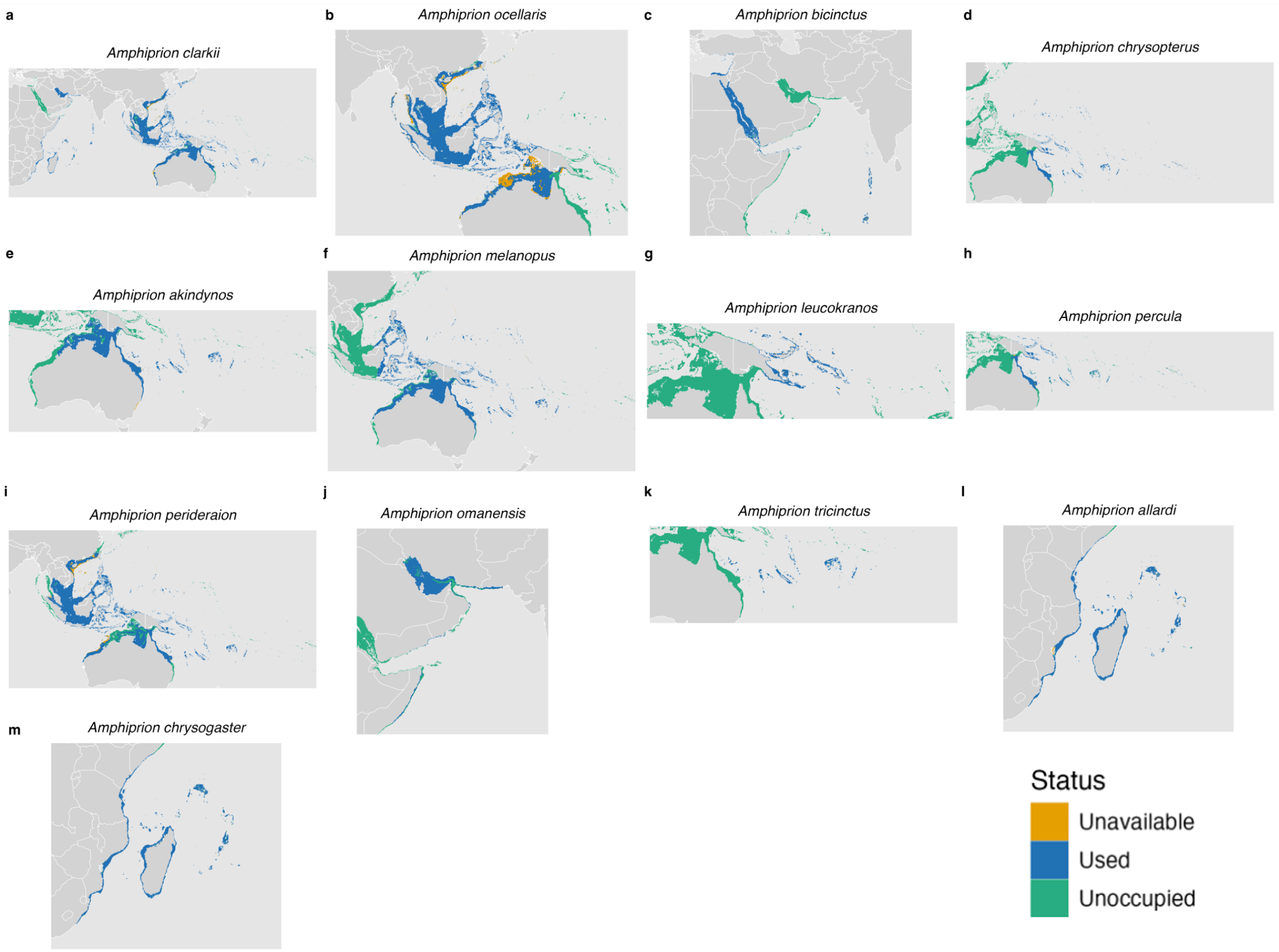


**Fig. S13**: **Spatial Distribution of Unavailable, Unoccupied, and Used (UUU) Proportions for Generalist Clownfish Species** Spatial patterns of environmental availability and utilization for 13 generalist clownfish species across their respective ranges in the Indo-Pacific region. Each panel represents a different species: a) *Amphiprion clarkii* b) *A. ocellaris* c) *A. bicinctus* d) *A. chrysopterus* e) *A. akindynos* f) *A. melanopus* g) *A. leucokranos* h) *A. percula* i) *A. perideraion* j) *A. omanensis* k) *A. tricinctus* l) *A. allardi* m) *A. chrysogaster* For each species, the maps display the proportions of Unavailable (areas suitable but inaccessible), Unoccupied (areas accessible but not utilized), and Used (areas both suitable and occupied) environments. Colors indicate the relative proportions of each UUU parameter, as detailed in the legend located at the bottom right. The variation in UUU patterns across species reflects differences in their ecological flexibility, host anemone distributions, and historical biogeography.

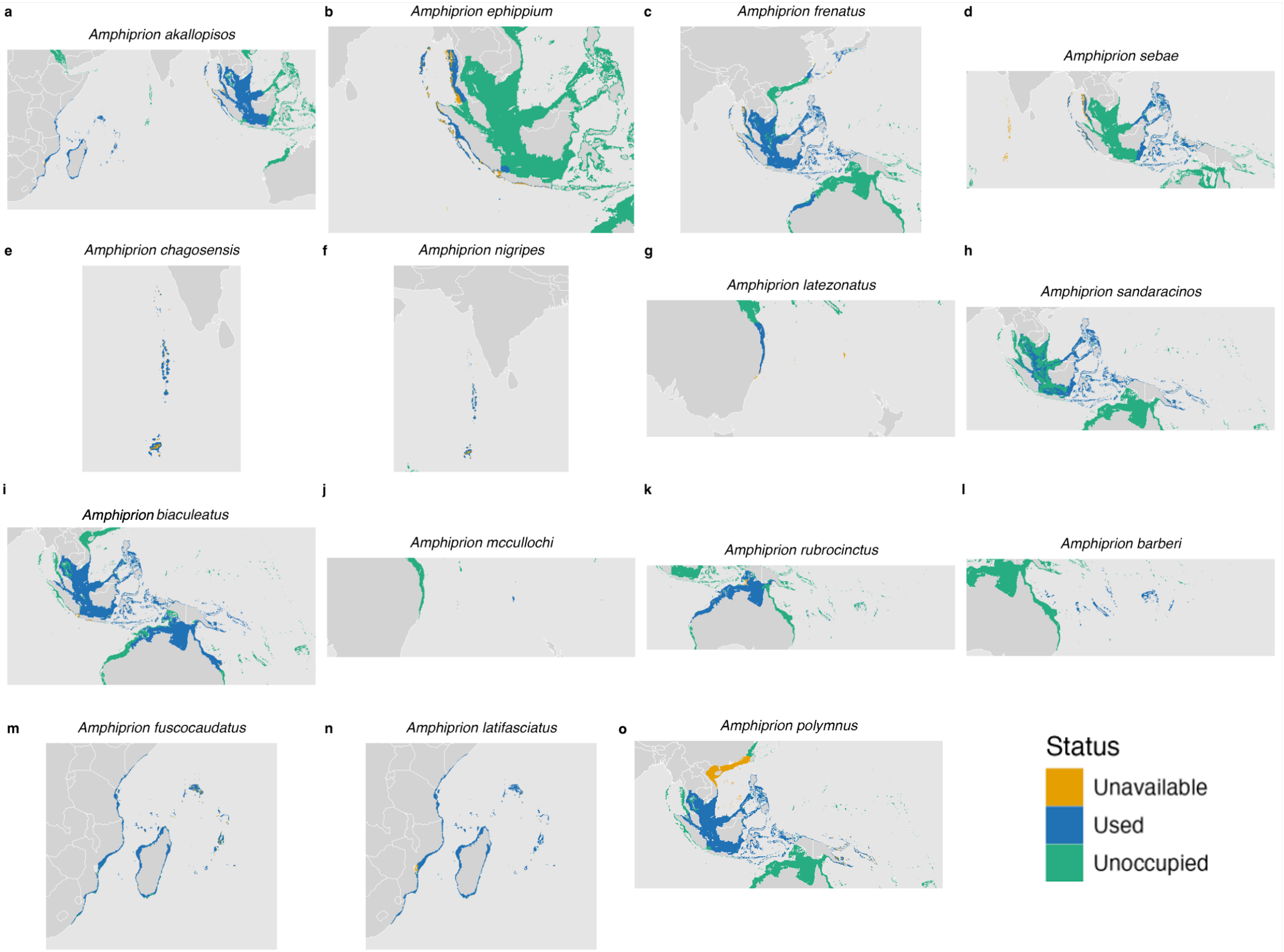


**Fig. S14**: **Spatial Distribution of Unavailable, Unoccupied, and Used (UUU) Proportions for Specialist Clownfish Species** Spatial patterns of environmental availability and utilization for 15 specialist clownfish species across their respective ranges in the Indo-Pacific region. Each panel corresponds to a different species: a) *Amphiprion akallopisos* b) *A. barberi* c) *A. chagosensis* d) *A. ephippium* e) *A. frenatus* f) *A. fuscocaudatus* g) *A. latezonatus* h) *A. latifasciatus* i) *A. mccullochi* j) *A. nigripes* k) *A. polymnus* l) *A. rubrocinctus* m) *A. sandaracinos* n) *A. sebae* o) *A. biaculeatus* For each species, the maps display the proportions of Unavailable (environmentally suitable but inaccessible), Unoccupied (accessible but not utilized), and Used (both suitable and occupied) areas. Colors indicate the relative proportions of each UUU parameter, as detailed in the legend located at the bottom right. These spatial UUU distributions are valuable for identifying potentially vulnerable or sink populations, as well as highlighting areas that may require increased sampling efforts.

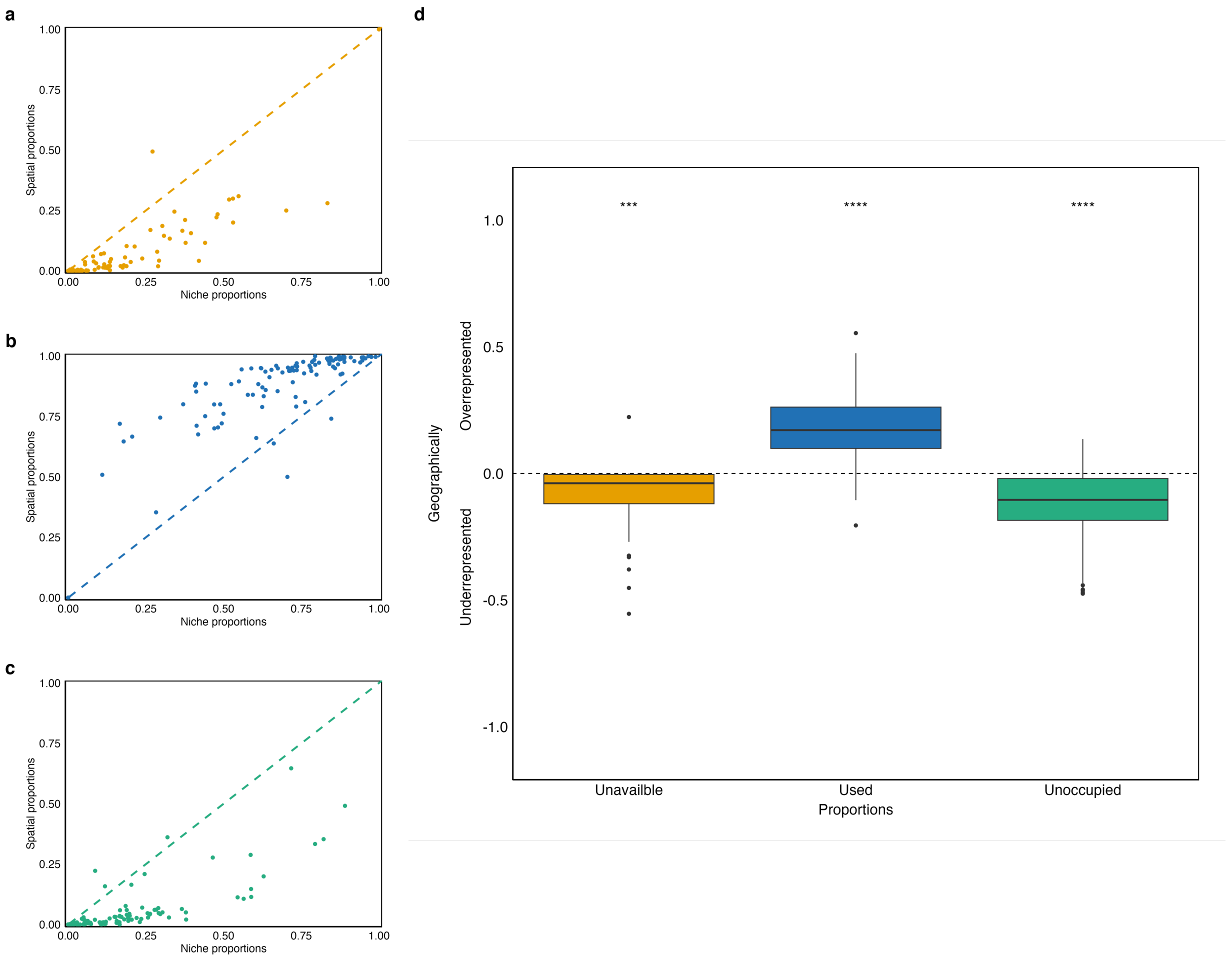


**Fig. S15**: **Comparison of Niche and Spatial UUU Proportions Across Marine Provinces** Relationship between niche-based and spatial-based Unavailable, Used, and Unoccupied (UUU) proportions for clownfish-anemone interactions across marine provinces. Panels a-c present scatterplots comparing niche-based proportions (x-axis) to their corresponding spatial representations (y-axis) for Unavailable, Used, and Unoccupied environments, respectively. In each of these panels, the colored dashed line represents a 1:1 relationship, indicating perfect equivalence between niche and spatial proportions. Panel d displays boxplots showing the distribution of differences between spatial and niche proportions for each UUU category: Unavailable (suitable but inaccessible), Used (suitable and accessible), and Unoccupied (accessible but unsuitable) environments. The dashed line at zero in panel d represents equal niche and spatial proportions. Asterisks above each boxplot indicate statistical significance of the difference from zero, with n.s. (not significant), * ($p < 0.05$), ** ($p < 0.01$), *** ($p < 0.001$), and **** ($p < 0.0001$). This analysis reveals discrepancies between environmental suitability (niche-based) and actual geographical distribution (spatial-based) of clownfish-anemone interactions.

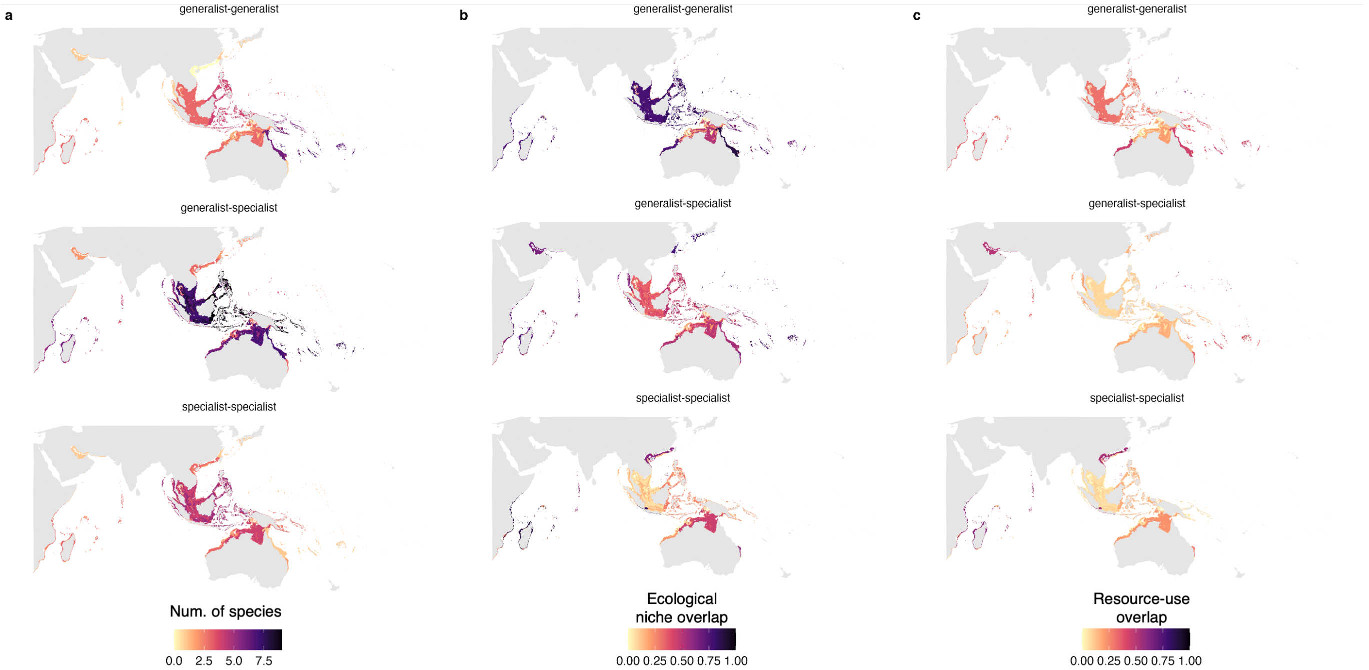


**Fig. S16**: **Geographical Patterns of Clownfish Species Interactions and Niche Overlap** Spatial distribution of clownfish species interactions and niche overlap across their range. Panel (a) depicts the geographical distribution of interaction types, showing the number of species involved in generalist-generalist, generalist-specialist, and specialist-specialist interactions at each location. The color intensity represents the species richness for each interaction type. Panel (b) displays the spatial distribution of averaged ecological niche overlap for each interaction type. Here, the color gradient indicates the magnitude of ecological nivche overlap, with darker colors representing higher overlap values. Panel (c) presents the spatial distribution of averaged resource-use overlap for each interaction type, again using a color gradient to show the intensity of overlap. This multi-faceted visualization allows for the comparison of species interaction patterns with both ecological and resource-use niche overlap across geographic space. The figure reveals potential hotspots of species interactions and areas of high overlap.

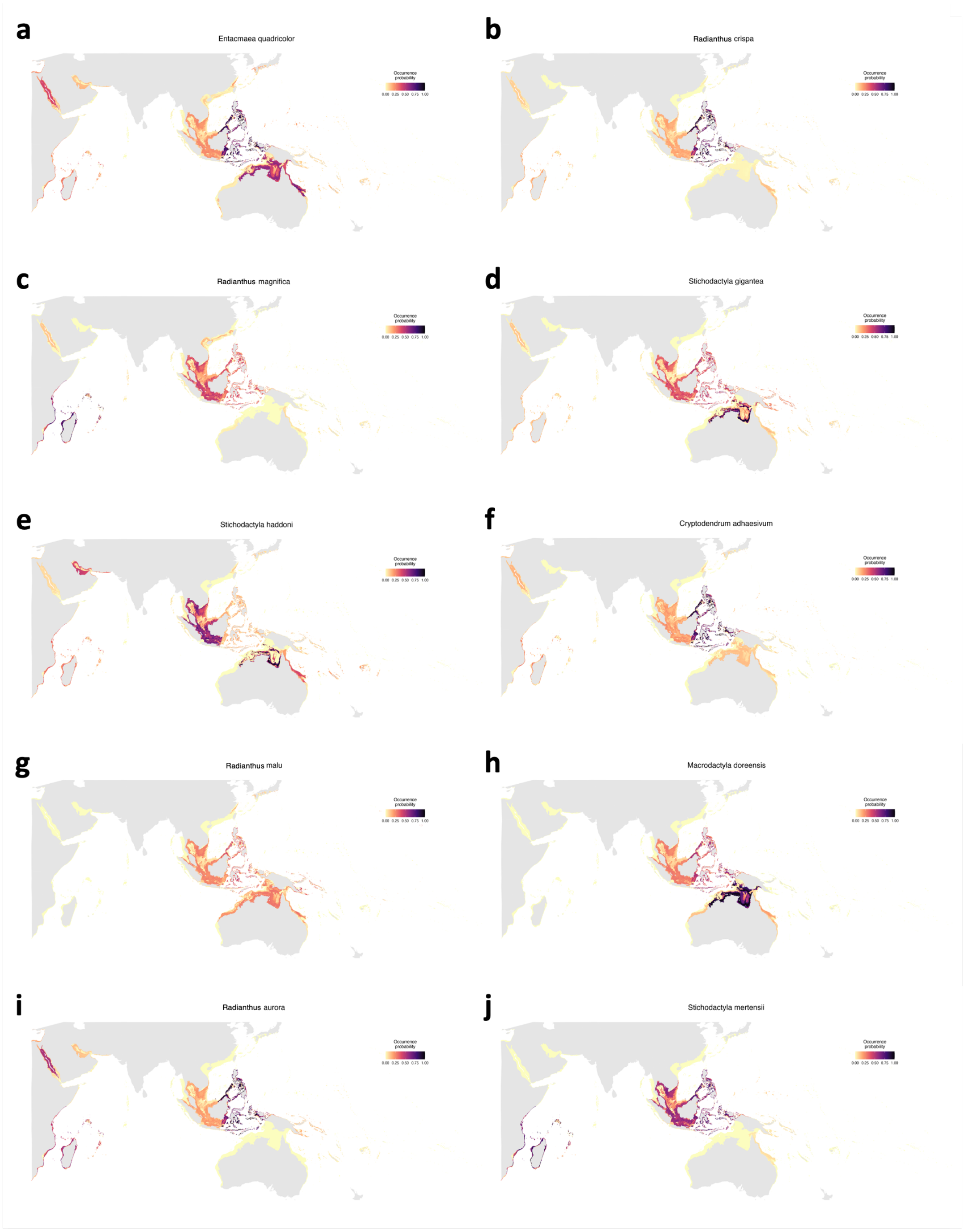


**Fig. S17**: **Geographical Distribution of Host Sea Anemone Species for Clownfish** Predicted distribution maps for ten sea anemone species that serve as hosts for clownfish, based on species distribution modeling. Each panel represents a different species: a) *Entacmaea quadricolor* b) *Radianthus crispa* (formerly *Heteractis crispa*) c) *Radianthus magnifica* (formerly *Heteractis magnifica*) d) *Stichodactyla gigantea* e) *Stichodactyla haddoni* f) *Cryptodendrum adhaesivum* g) *Radianthus malu* (formerly *Heteractis malu*) h) *Macrodactyla doreensis* i) *Radianthus aurora* (formerly *Heteractis aurora*) j) *Stichodactyla mertensii* The maps illustrate the probability of occurrence presence of each species across the Indo-Pacific region. Darker colors indicate a higher probability of occurrence, while lighter colors represent lower probabilities and areas of absence.

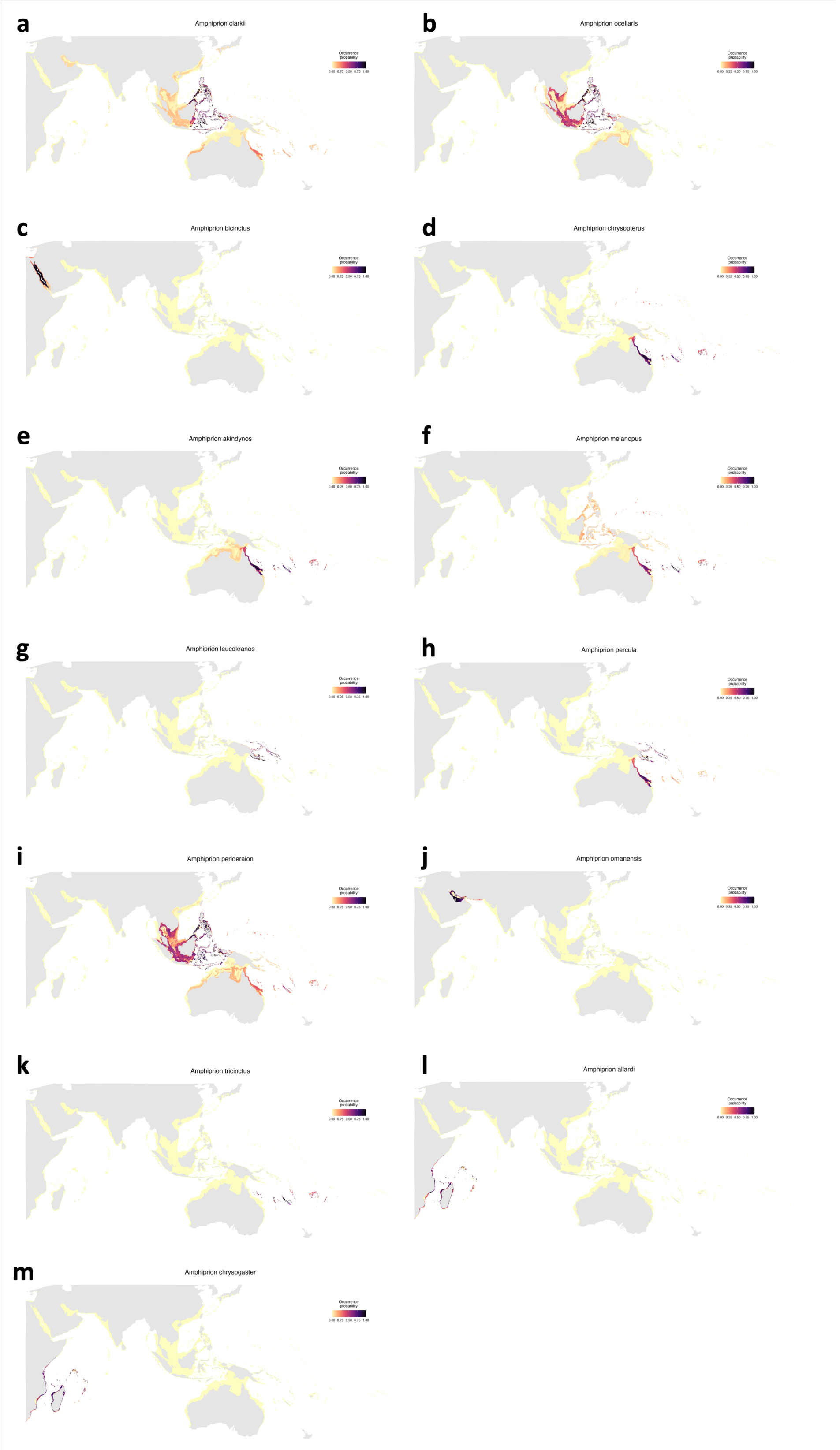


**Fig. S18**: **Predicted Geographical Distributions of Generalist Clownfish Species** Distribution maps for thirteen generalist clownfish species of the genus *Amphiprion*. Each panel corresponds to a different species: a) *A. clarkii* b) *A. ocellaris* c) *A. bicinctus* d) *A. chrysopterus* e) *A. akindynos* f) *A. melanopus* g) *A. leucokranos* h) *A. percula* i) *A. perideraion* j) *A. omanensis* k) *A. tricinctus* l) *A. allardi* m) *A. chrysogaster* The maps illustrate the probability of occurrence presence of each species across the Indo-Pacific region. Darker colors indicate a higher probability of occurrence, while lighter colors represent lower probabilities and areas of absence. These distributions are based on species distribution models that incorporate both environmental variables and host anemone availability.

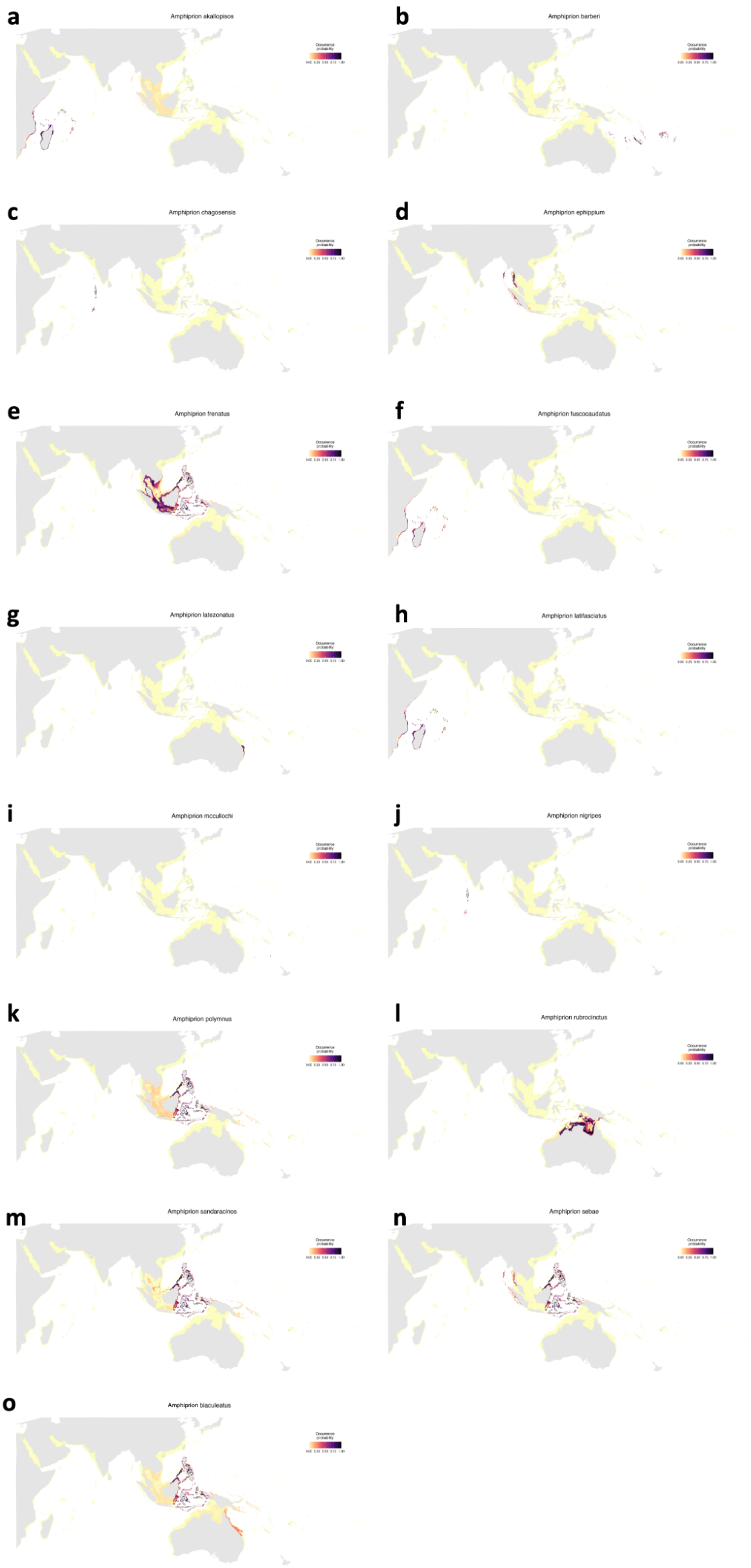


**Fig. S19**: **Predicted Geographical Distributions of Specialist Clownfish Species** Distribution maps for fifteen specialist clownfish species of the genus *Amphiprion*: a) *A. akallopisos* b) *A. ephippium* c) *A. frenatus* d) *A. sebae* e) *A. chagosensis* f) *A. nigripes* g) *A. latezonatus* h) *A. sandaracinos* i) *A. biaculeatus* j) *A. mccullochi* k) *A. rubrocinctus* l) *A. barberi* m) *A. fuscocaudatus* n) *A. latifasciatus* o) *A. polymnus* The maps illustrate the probability of occurrence presence of each species across the Indo-Pacific region. Darker colors indicate a higher probability of occurrence, while lighter colors represent lower probabilities and areas of absence. These distributions are derived from species distribution models incorporating both environmental variables and the availability of specific host anemone species.

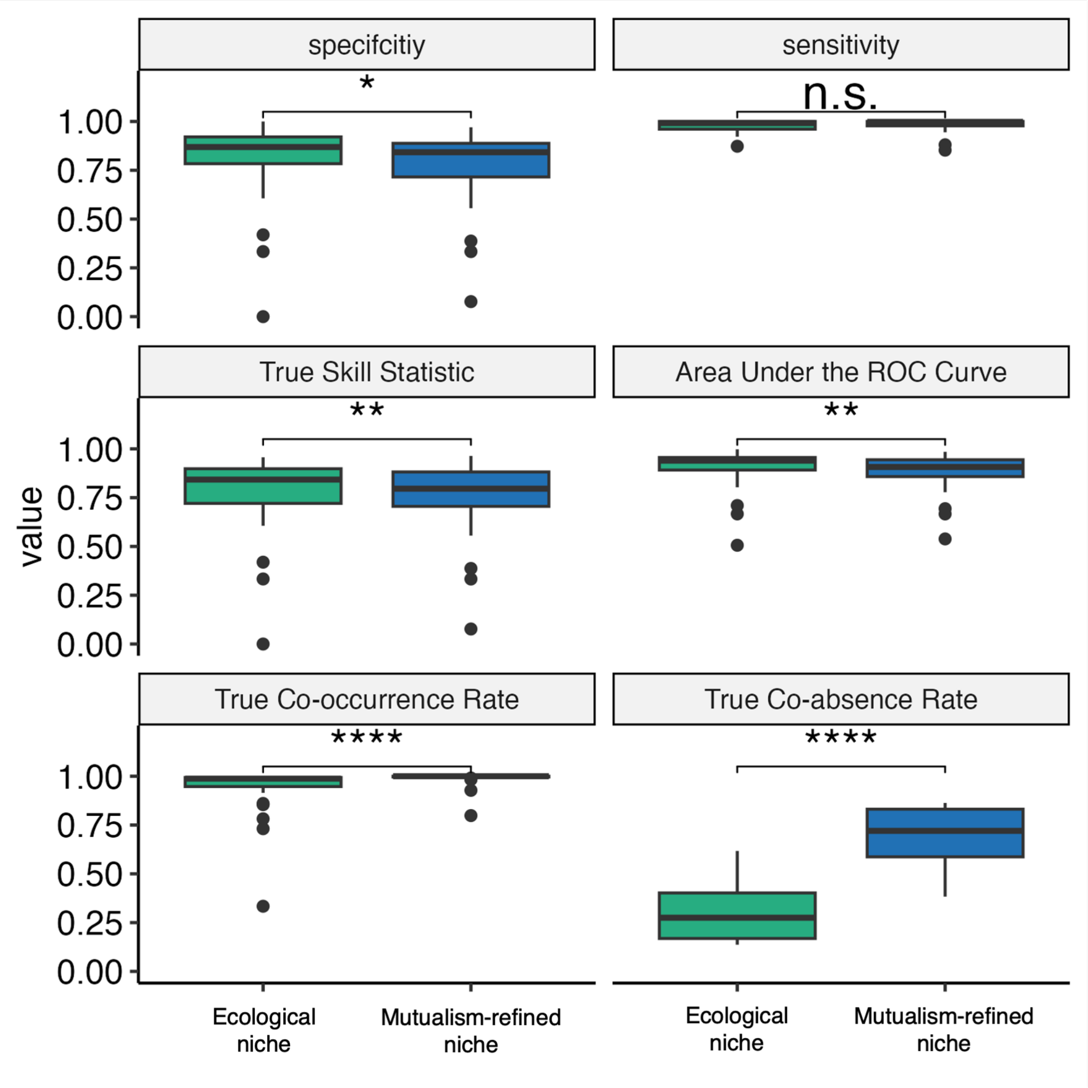


**Fig. S20**: **Comparative Evaluation of Ecological and Mutualism-Refined Niche Models for Clownfish Species** Key evaluation metrics for two types of species distribution models: Ecological Niche and Mutualism-Refined models, applied to 28 clownfish species. The boxplots display the median and Interquartile Range (IQR) for each metric across all modeled species. Two critical metrics are shown at the bottom: (1) True Co-occurrence Rate, which quantifies the proportion of locations where both the focal clownfish and at least one of its host anemones are predicted to be present, relative to the total predicted presences of the clownfish; and (2) True Co-Absence Rate, representing the proportion of locations where both the focal clownfish and all its potential host anemones are predicted to be absent, relative to the total predicted absences of the clownfish. These metrics assess the models' ability to capture the obligate mutualistic relationship between clownfish and their host anemones. Statistical comparisons between Ecological Niche and Mutualism-Refined models were conducted using Wilcoxon's paired Rank test, with significance levels indicated as: n.s. (not significant), * ($p < 0.05$), ** ($p < 0.01$), *** ($p < 0.001$), and **** ($p < 0.0001$).

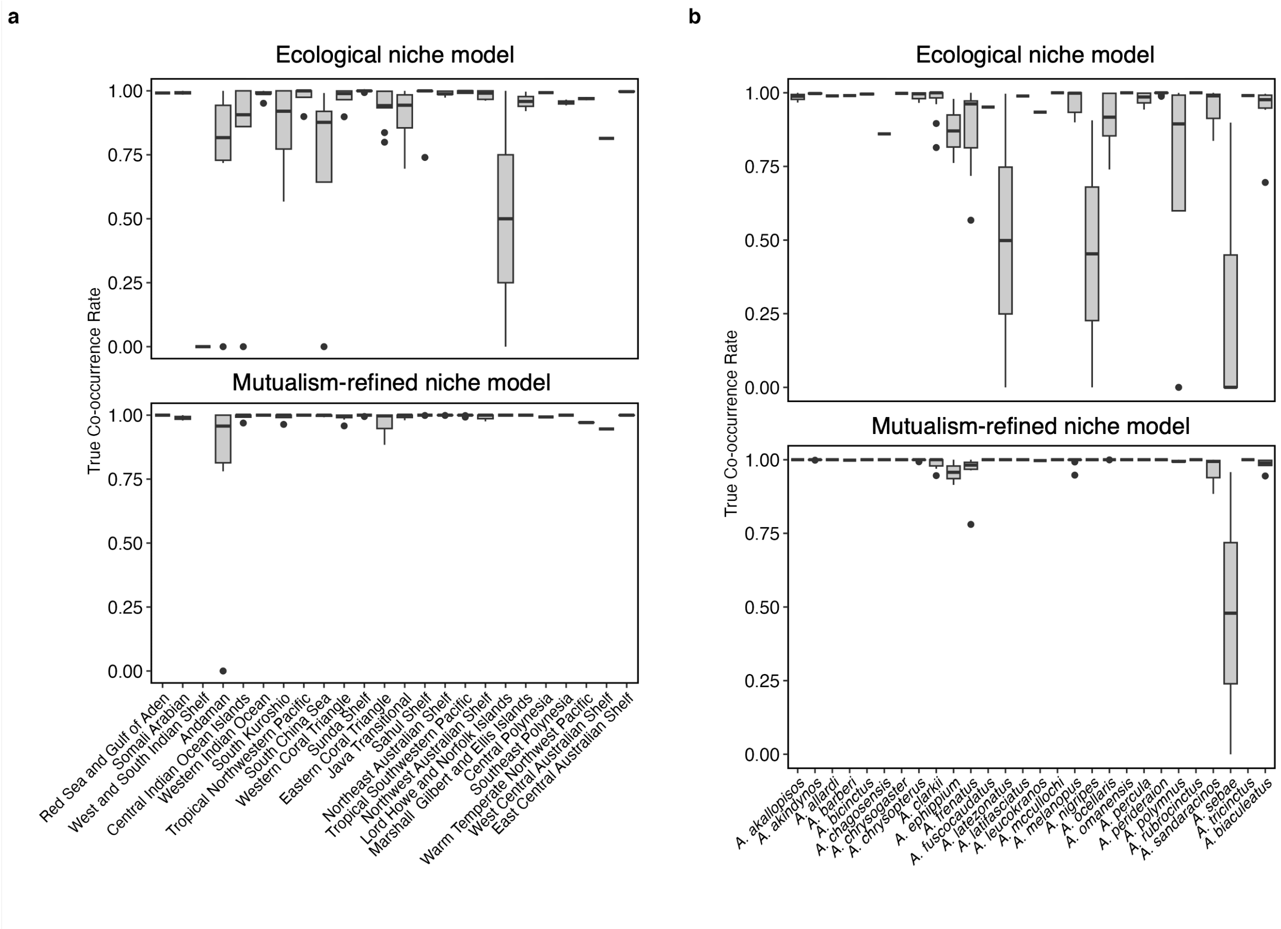


**Fig. S21**: **Comparative Analysis of True Co-occurrence Rates in Ecological and Mutualism-Refined Niche Models** Comparison of True Co-occurrence Rates between Ecological Niche and Mutualism-Refined models for clownfish-anemone symbioses, analyzed across both geographic and taxonomic dimensions. Panel (a) illustrates the variation in True Co-occurrence Rate across 27 marine provinces for each model type. Here, each boxplot represents the median and Interquartile Range (IQR) of rates observed within a province, encompassing all clownfish species present. This geographical perspective reveals how the accuracy of predicting clownfish-anemone co-occurrences varies across different marine regions. Panel (b) displays the True Co-occurrence Rate for each of the 28 modeled clownfish species, comparing EN and BC models. In this panel, each boxplot represents the median and IQR of rates observed for a single species across its entire range. This species-specific view highlights potential differences in model performance that may be related to individual species' ecological traits or host specificity. True Co-occurrence Rate, defined as the proportion of locations where both clownfish and at least one of its host anemones are predicted to be present relative to the total predicted presences of the clownfish, serves as a critical metric for assessing model accuracy in capturing this obligate mutualism.

### Supplementary Material & Methods

#### Generalizing the Framework to Other Types of Biotic Interactions

The framework developed in this study can be extended to encompass a broader range of biotic interactions, both positive and negative. The interaction matrix can accommodate values from [-1, 1], representing a continuous gradient of biotic effects. A value of -1 indicates a strictly negative interaction, where the presence of the biotic interactor precludes the focal species if the interactor's relative abundance equals or exceeds that of the focal species. A value of 0 represents a neutral interaction, while a value of 1 signifies a strictly positive interaction, as demonstrated in this study.

In this generalized framework, the multi-species niche (referred to as host availability in our study) represents ecological availability in terms of restriction or facilitation by biotic interactions. We propose the following equation to estimate ecological availability:

$$\omega = \left(1 - \prod_{k=1}^{p} (1 - o_k \alpha_k)\right) \prod_{k=1}^{n} (1 - o_k \alpha_k)$$

where the first term of the equation represents the probability of positive interactions occurring and the second term represents the probability of negative interactions being absent. The value of □ indicates the availability score that reflects ecological suitability for the focal species concerning biotic interactions.

Subsequently, we can calculate the mutualism-refined occurrence density $o'$ of the focal species, considering biotic interactions:

$$o' = (1 - I)o + I\omega o$$

where $I$ ranges from 0 to 1 and represents the degree of dependence of the focal species' niche on existing biotic interactions. The original occurrence density is denoted by $o$. This generalized framework allows for a more comprehensive analysis of species distributions by incorporating the nuanced effects of various biotic interactions across ecological systems.

#### Species Distributions

To infer species distributions (Fig. S17, S18 and S19), we projected both the occurrence density ($o$) and the mutualism-refined occurrence density ($o'$) of each environment estimated for all provinces onto their corresponding geographical coordinates. For each species, we assigned suitability values to specific locations based on the occurrence density of the environment in the niche space. We evaluated the predictive performance of each model using the True Skill Statistic (TSS), which combines specificity (True Negative Rate) and sensitivity (True Positive Rate) (Figure S20). To facilitate this evaluation, we generated pseudo-absences using a random stratified selection process. This process employed a vector of probabilities derived from the average distances to true occurrences within the environmental envelope, with greater distances indicating a higher probability of absence. This approach ensures a more realistic representation of potential species absences across the environmental gradient.

To transform the continuous species suitability distributions into binary presence/absence predictions, we employed the maxTSS approach (Guisan *et al.* 2017). This method selects the suitability threshold that maximizes the TSS across all possible thresholds, optimizing the balance between sensitivity and specificity. This comprehensive approach to modeling and evaluating species distributions allows us to capture both the fundamental niche requirements and the influence of mutualistic interactions on realized species distributions. By comparing the outputs of models based on $o$ and $o'$, we assessed the impact of incorporating host availability on the predicted distributions of clownfish species across their range (Fig. S20 and S21.

#### Niche Metrics

**Niche UUU Proportions:** We adapted the COUE terminology (Guisan *et al.*, 2014) to analyze the effect of mutualistic interactions on clownfish niches, drawing an analogy with niche shifts during invasions. In our framework, 'Unavailable' refers to the portion of the clownfish ecological niche inaccessible due to host absence. 'Unoccupied' denotes the portion of environmental availability unsuitable for the focal species, representing potential niche expansion areas. 'Used' represents the effective niche over the union of ecological niche and environmental availability.

**Centroid Shift:** We quantified the shift in niche position between the ecological and mutualism-refined niches for each species and province using Euclidean distance between the two niche centroids.

**Environmental Shift:** We measured the angle between environmental and mutualism-refined niche centroids relative to a complete inversion (180° rotation) using the `angle.calc` function from the *Morpho* R package (Schlager, 2017). This shift represents the change in environmental variable weights in the mutualism-refined niche due to host availability.

**Niche Dissimilarity:** We employed Schoener's $D$ metric (Schoener, 1970; Warren et al., 2008) to measure niche overlap between ecological and mutualism-refined niches ($D_{ecological-mutualism\text{-}refined}$). We calculated $1 -$

$D_{ecological-mutualism\text{-}refined}$ to estimate the niche shift caused by mutualistic interactions, with higher values indicating stronger host availability effects on the clownfish niche.

#### Spatial Metrics

**Spatial UUU Parameters:** We applied UUU parameters to geographical distributions. 'Unavailable' represents the portion of the species range inaccessible due to lack of mutualistic interactions. 'Unoccupied' denotes the portion of the hosts' range unsuitable for clownfish but potentially suitable through mutualistic facilitation. 'Used' represents the stable portion of the species range fitting both environmental suitability and mutualistic accessibility.

#### Resource-Use Overlap

To estimate a more realistic competition proxy, we developed a method that considers host use as a primary resource (Fig. S2). For each pairwise estimate, we replicated the environmental space into layers equal to the number of shared hosts. We then replicated species niches onto layers corresponding to their hosts, computed niche overlap on each layer, and averaged the estimates. This approach yields higher competition values for species with common hosts and shared environments, decreasing with the number of alternative hosts

#### Sensitivity Analysis

We conducted sensitivity analyses to assess different association matrices' effects on the framework. We ran the framework on Environmental Niche models using matrices ranging from unconstrained mutualism to maximized mutualistic constraint. The models are described in Table S6.

We assessed the effects on niche unavailability, spatial unavailability, proportion of hosts used relative to occurring sea anemones per region, and resource-use overlap in clownfish species at a regional level.

### Supplementary Results & Discussion

Sensitivity analyses confirm the relationship between interaction numbers and niche/distribution constraints (Fig. S4 A and B), as well as system complexity and resource overlap. The observed system (known clownfish-host interactions) shows a lower proportion of hosts used per region per species than randomized models, suggesting a resource usage maximization strategy (Fig. S4 C).

Fig. S4 D shows a similar response between model `rRrGoN` and the `Observed model` indicating that the overall structure of clownfish-anemone associations is largely determined by the number of interactions available, rather than the specific identity of the host species involved. The higher resource overlap observed in the `rRrGoN` and `Observed` models compared to other randomized models likely reflects the evolutionary history and phylogenetic constraints of clownfish-anemone associations. This pattern suggests that host specificity in closely related clownfish species is not merely a product of random associations but rather the result of shared evolutionary trajectories and conserved traits like mutualistic behavior. The persistence of these patterns, even when hosts are randomized within the constraints of interaction numbers, indicates that closely related clownfish species tend to exhibit similar host preferences, possibly due to shared physiological or behavioral adaptations. Additionally, the number and distribution of host associations may be influenced by historical biogeographical factors that have shaped clownfish diversification. This finding underscores the importance of considering phylogenetic relationships and evolutionary history when interpreting patterns of host use and resource overlap in symbiotic systems like the clownfish-anemone mutualism. Overall, it suggests that the observed patterns of host specificity may represent adaptive solutions shaped by ecological interactions and evolutionary constraints over time.
